# Robust Estimation of Bacterial Cell Count from Optical Density

**DOI:** 10.1101/803239

**Authors:** Jacob Beal, Natalie G. Farny, Traci Haddock-Angelli, Vinoo Selvarajah, Geoff S. Baldwin, Russell Buckley-Taylor, Markus Gershater, Daisuke Kiga, John Marken, Vishal Sanchania, Abigail Sison, Christopher T. Workman, the iGEM Interlab Study Contributors

**Author notes:** Membership list is provided in Supplementary Note: iGEM Interlab Study Contributors. (J.B.), (N.G.F.), (T.H-A.), (G.S.B.), (M.G.), (C.T.W.).

## Abstract

Optical density (OD) is a fast, cheap, and high-throughput measurement widely used to estimate the density of cells in liquid culture. These measurements, however, cannot be compared between instruments without a standardized calibration protocol and are challenging to relate to actual cell count. We address these shortcomings with an interlaboratory study comparing three OD calibration protocols, as applied to eight strains of *E. coli* engineered to constitutively express varying levels of GFP. These three protocols—comparison with colloidal silica (LUDOX), serial dilution of silica microspheres, and a reference colony-forming unit (CFU) assay—are all simple, low-cost, and highly accessible. Based on the results produced by the 244 teams completing this interlaboratory study, we recommend calibrating OD using serial dilution of silica microspheres, which readily produces highly precise calibration (95.5% of teams having residuals less than 1.2-fold), is easily assessed for quality control, and as a side effect also assesses the effective linear range of an instrument. Moreover, estimates of cell count from silica microspheres can be combined with fluorescence calibration against fluorescein to obtain units of Molecules of Equivalent Fluorescein (MEFL), allowing direct comparison and data fusion with equivalently calibrated flow cytometry measurements: in our study, fluorescence per cell measurements showed only a 1.07-fold mean difference between plate reader and flow cytometry data.

## Introduction

Comparable measurements are a *sine qua non* for both science and engineering, and one of the most commonly needed measurements of microbes is the number (or concentration) of cells in a sample. The most common method for estimating the number of cells in a liquid suspension is the use of optical density measurements (OD) at an absorbance wavelength of 600nm (OD600) [1]. The dominance of OD measurements is unsurprising, particularly in plate readers, as these measurements are extremely fast, inexpensive, simple, relatively non-disruptive, high-throughput, and readily automated. Alternative measurements of cell count—microscopy (with or without hemocytometer), flow cytometry, colony forming units (CFU), and others, e.g., [2–5]—lack many of these properties, though some offer other benefits, such as distinguishing viability and being unaffected by cell states such as inclusion body formation, protein expression, or filamentous growth [6].

A key shortcoming of OD measurements is that they do not actually provide a direct measure of cell count. Indeed, OD is not even linearly related to cell count except within a limited range [7]. Furthermore, because the phenomenon is based on light scatter rather than absorbance, it is relative to the configuration of a particular instrument. Thus, in order to relate OD measurements to cell count—or even just to compare measurements between instruments and experiments—it is necessary to establish a calibration protocol, such as comparison to a reference material.

While the problems of interpreting OD values have been studied (e.g., [1, 6, 7]), no previous study has attempted to establish a standard protocol to reliably calibrate estimation of cell count from OD. To assess reliability, it is desirable to involve a large diversity of instruments and laboratories, such as those participating in the International Genetically Engineered Machines (iGEM) competition [8], where hundreds of teams at the high school, undergraduate, and graduate levels been organized previously to study reproducibility and calibration for fluorescence measurements in engineered *E. coli* [9, 10]. As iGEM teams have a high variability in training and available resources, organizing an interlaboratory study with iGEM also demands that protocols be simple, low cost, and highly accessible. The large scale and high variability between teams also allows investigation of protocol robustness, as well as how readily issues can be identified and debugged in protocol execution.

We thus organized a large-scale interlaboratory study within iGEM to compare three candidate OD calibration protocols: a colony-forming unit (CFU) assay, the de facto standard assay for determining viable cell count; comparison with colloidal silica (LUDOX) and water, previously used for normalizing fluorescence measurements [9]; and serial dilution of silica microspheres, a new protocol based on a recent study of microbial growth [7]. Overall, this study demonstrates that serial dilution of silica microspheres is by far the best of these three protocols, allowing highly precise, accurate, and robust calibration that is easily assessed for quality control and can also evaluate the effective linear range of an instrument.

## Results

To evaluate the three candidate OD calibration protocols, we organized an interlaboratory study as part of the 2018 International Genetically Engineered Machine (iGEM) competition. The precision and robustness of each protocol is assessed based on the variability between replicates, between reference levels, and between laboratories. The overall efficacy of the protocols was then further evaluated based on the reproducibility of cross-laboratory measurements of cellular fluorescence, as normalized by calibrated OD measurements.

### Experimental Data Collection

Each contributing team was provided with a set of calibration materials and a collection of eight engineered genetic constructs for constitutive expression of GFP at a variety of levels. Specifically, the constructs consisted of a negative control, a positive control, and six test constructs that were identical except for promoters from the Anderson library [11], selected to give a range of GFP expression (illustrated in Figure 1(a), with complete details provided in Supplementary Data 1 DNA Constructs). These materials were then used to follow a calibration and cell measurement protocol (Materials and Methods; Supplementary Note: Plate Reader and CFU Protocol and Supplementary Note: Flow Cytometer Protocol).

**Fig 1.**
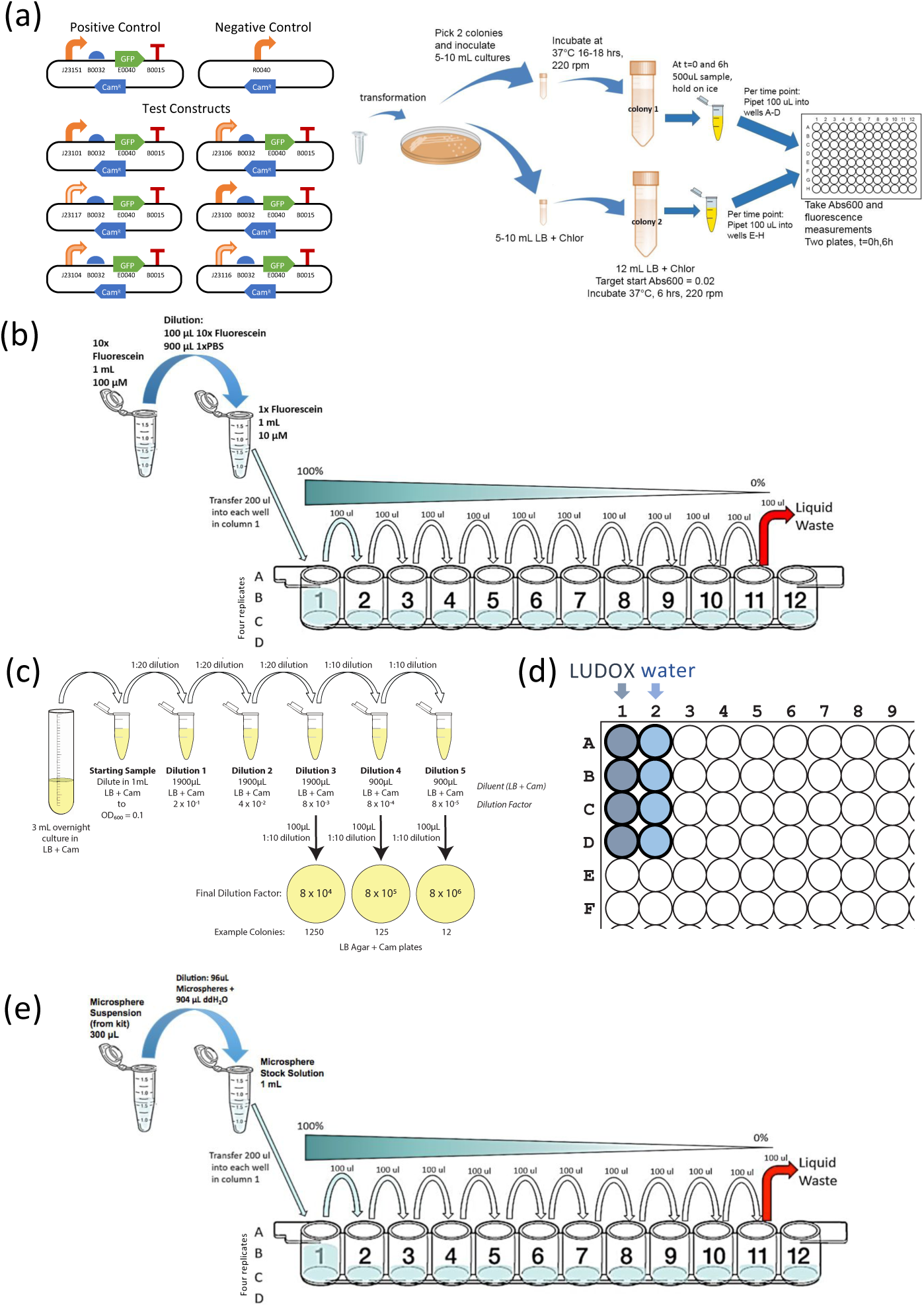
Study design: (a) each team cultured eight strains of engineered E. coli expressing GFP at various levels: positive and negative controls plus a library of six test constructs with promoters selected to give a range of levels of expression. Each team also collected four sets of calibration measurements, (b) fluorescein titration for calibration of GFP fluorescence, plus three alternative protocols for calibration of absorbance at 600nm: (c) dilution and growth for colony forming units (CFU), (d) LUDOX and water, and (e) serial dilution of 0.961*µ*m diameter monodisperse silica microspheres.

Each team transformed *E. coli* K-12 DH5-alpha with the provided genetic constructs, culturing two biological replicates for each of the eight constructs. Teams measured absorbance at 600nm (OD600) and GFP in a plate reader from 4 technical replicates per biological replicate at the 0 and 6 hour time points, along with media blanks, thus producing a total of 144 OD600 and 144 GFP measurements per team. Teams with access to a flow cytometer were asked to also collect GFP and scatter measurements for each sample, plus a sample of SpheroTech Rainbow Calibration Beads [12] for fluorescence calibration.

Measurements of GFP fluorescence were calibrated using serial dilution of fluorescein with PBS in quadruplicate, using the protocol from [9], as illustrated in Figure 1(b). Starting with a known concentration of fluorescein in PBS means that there is a known number of fluorescein molecules per well. The number of molecules per arbitrary fluorescence unit can then by estimated by dividing the expected number of molecules in each well by the measured fluorescence for the well; a similar computation can be made for concentration.

Measurements of OD via absorbance at 600nm (OD600) were calibrated using three protocols and for each of these a model was devised for the purpose of fitting the data obtained in the study (Methods):

- Calibration to colony forming units (CFU), illustrated in Figure 1(c): four overnight cultures (two each of positive and negative controls), were sampled in triplicate, each sample diluted to 0.1 OD, then serially diluted, and the final three dilutions spread onto bacterial culture plates for incubation and colony counting (a total of 36 plates per team). The number of CFU per OD per mL is estimated by multiplying colony count by dilution multiple. This protocol has the advantage of being well established and insensitive to non-viable cells and debris, but the disadvantages of an unclear number of cells per CFU, potentially high statistical variability when the number of colonies is low, and being labor intensive.
- Comparison of colloidal silica (LUDOX CL-X) and water, illustrated in Figure 1(d): this protocol is adapted from [9] by substitution of a colloidal silica formulation that is more dense and freeze-tolerant (for easier shipping). Quadruplicate measurements are made for both LUDOX CL-X and water, with conversion from arbitrary units to OD measurement in a standard spectrophotometer cuvette estimated as the ratio of their difference to the OD measurement for LUDOX CL-X in a reference spectrophotometer. This protocol has the advantage of using extremely cheap and stable materials, but the disadvantage that LUDOX CL-X provides only a single reference value, and that it calibrates for instrument differences in determination of OD but cannot determine the number of particles.
- Comparison with serial dilution of silica microspheres, illustrated in Figure 1(e). This novel protocol, inspired by the relationship between particle size, count, and OD [7], uses quadruplicate serial dilution protocol of 0.961*µ*m diameter monodisperse silica microspheres (selected to match the approximate volume and optical properties of *E. coli*) in water (similar to fluorescein dilution, but with different materials). With a known starting concentration of particles, the number of particles per OD600 unit is estimated by dividing the expected number of particles in each well by the measured OD for the well. This protocol has the advantages of low cost and of directly mapping between particles and OD, but the disadvantage that the microspheres tend to settle and are freeze-sensitive.

Data from each team were accepted only if they met a set of minimal data quality criteria (Supplementary Note: Data Acceptance Criteria), including values being non-negative, the positive control being significantly brighter than the negative control, and measured values for calibrants decreasing as dilution increases. In total, 244 teams provided data meeting these minimal criteria, with 17 teams also providing usable flow cytometry data. Complete anonymized data sets and analysis results are available in Supplementary Data 2 Complete Data.

### Robustness of calibration protocols

We assessed the robustness of the calibration protocols under test in two ways: replicate precision and residuals. Replicate precision can be evaluated simply in terms of the similarity of values for each technical replicate of a protocol. The smaller the coefficient of variation (i.e., ratio of standard deviation to mean), the more precise the protocol. With regards to residuals, on the other hand, we considered the modeled mechanism that underlies each calibration method and assess how well it fits the data. Here, the residual is the distance between each measured value provided by a team and the predicted value of a model fit using that same set of data (see Materials and Methods for details of each mechanism model and residual calculations). The smaller the residual value, the more precise the protocol. Moreover, the more similar the replicate precision and residuals across teams, the more robust the protocol is to variations in execution conditions.

Figure 2 shows the distribution of the coefficients of variation (CVs) for all valid replicates for each of the calibrant materials (see Materials and Methods for validity criteria). For CFU, basic sampling theory implies that the dilution with the largest number of countably distinct colonies (lowest dilution) should have the best CV, and indeed this is the case for 81.6% of the samples. This percentage is surprisingly low, however, and indicates a higher degree of variation than can be explained by the inherent stochasticity of the protocol: CFU sampling should follow a binomial distribution and have a little over 3-fold higher CV with each 10-fold dilution, but on average it was much less. This indicates the presence of a large component of variation with an unknown source, which is further confirmed by the fact that even the best CVs are quite high: the best of the three dilutions for each team has CV ≤ 0.1 for only 2.1% of all data sets and CV ≤ 0.2 for only 16.4% of all data sets.

**Fig 2.**
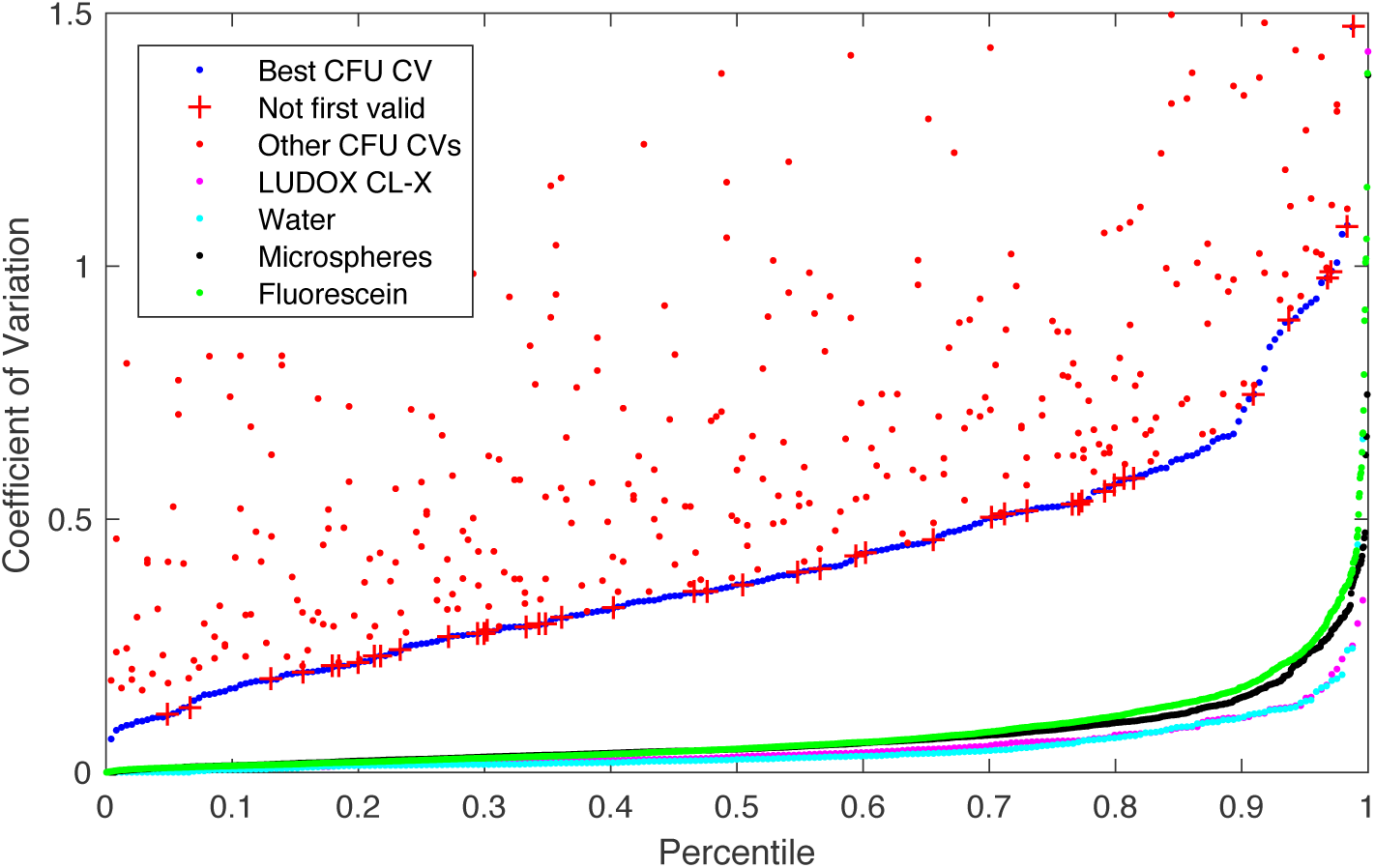
Distribution of the coefficient of variation for valid replicate sets in CFU, LUDOX/water, microspheres, and fluorescein. CFU models are generated from only the best CV dilution (blue); other dilutions are shown separately above. Even the best CV CFU dilutions, however, have a distribution far worse than the other four methods, and are surprisingly often not the lowest dilution (red crosses). Of the others, LUDOX (magenta) and water (light blue) have the best and near-identical distributions, while microspheres (black) and fluorescein (green) are only slightly higher.

LUDOX and water have the lowest CV, at CV ≤ 0.1 for 86.9% (LUDOX) and 88.1% (water) of all replicate sets and CV ≤ 0.2 for 97.1% (LUDOX) and 98.0% (water) of all replicate sets. Microspheres and fluorescein have slightly higher CV, at CV ≤ 0.1 for 80.8% (microspheres) and 76.9% (fluorescein) of all replicate sets and CV ≤ 0.2 for 93.9% (microspheres) and 92.4% (fluorescein) of all replicate sets. The difference between these two pairs likely derives from the fact that the LUDOX and water samples are each produced in only a single step, while the serial dilution of microspheres and fluorescein allows inaccuracies to compound in the production of later samples.

The accuracy of a calibration protocol is ultimately determined by how replicate data sets across the study are jointly interpreted to parameterize a model of the calibration protocol, one part of which is the scaling function that maps between arbitrary units and calibrated units. As noted above, this can be assessed by considering the residuals in the fit between observed values and their fit to the protocol model. To do this, we first estimated the calibration parameters from the observed experimental values (see Materials and Methods for the unit scaling computation for each calibration method), then used the resulting model to “predict” what those values should have been (e.g., 10-fold less colonies after a 10-fold dilution). The closer the ratio was to one, the more the protocol was operating in conformance with the theory supporting its use for calibration, and thus the more likely that the calibration process produced an accurate value.

Here we see a critical weakness of the LUDOX/water protocol: the LUDOX and water samples provide only two measurements, from which two model parameters are set: the background to subtract (set by water) and the scaling between background-subtracted LUDOX and the reference OD. Thus, the dimensionality of the model precisely matches the dimensionality of the experimental samples, and there are no residuals to assess. As such, the LUDOX/water protocol may indeed be accurate, but its accuracy cannot be empirically assessed from the data it produces. If anything goes wrong in the reagents, protocol execution, or instrument, such problems cannot be detected unless they are so great as to render the data clearly invalid (e.g., the OD of water being less than the OD of LUDOX).

The CFU protocol and the two serial dilution protocols, however, both have multiple dilution levels, overconstraining the model and allowing likely accuracy to be assessed. Figure 3 shows the distribution of residuals for these three protocols, in the form of a ratio between the observed mean for each replicate set and the value predicted by the model fit across all replicate sets. The CFU protocol again performs extremely poorly, as we might expect based on the poor CV of even the best replicates: only 7.3% of valid replicate sets have a residual within 1.1-fold, only 14.0% within 1.2-fold, and overall the geometric standard deviation of the residuals is 3.06-fold—meaning that values are only reliable to within approximately two orders of magnitude! Furthermore, the distribution is asymmetric, suggesting that the CFU protocol may be systematically underestimating the number of cells in the original sample. The accuracy of the CFU protocol thus appears highly unreliable.

**Fig 3.**
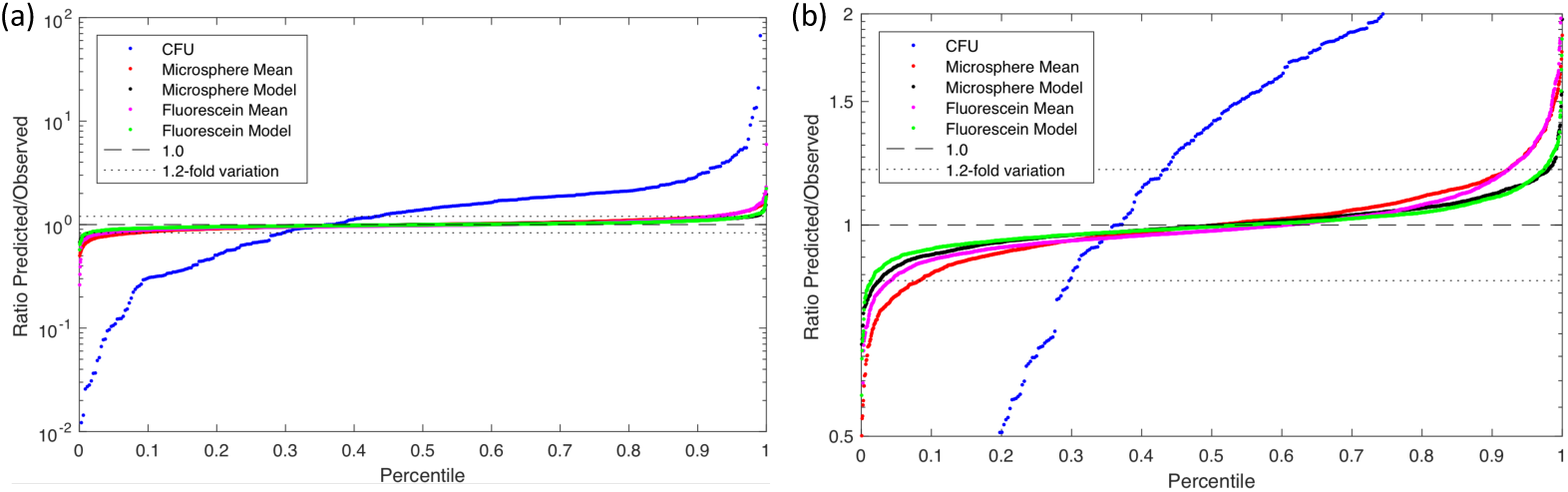
(a) Model fit residual distribution for each replica set in the CFU (blue), microsphere, and fluorescein calibration protocols. (b) Expanding the Y axis to focus on the microsphere and fluorescein distributions shows that incorporating a model parameter for systematic pipetting error (black, green) produces a significantly better fit (and thus likely more accurate unit calibration) than a simple geometric mean over scaling factors (red, magenta).

The microsphere dilution protocol, on the other hand, produced much more accurate results. Even with only a simple model of perfect dilution, the residuals are quite low (red line in Figure 3(b)), having 61.0% of valid replicates within 1.1-fold, 83.6% within 1.2-fold, and an overall geometric standard deviation of 1.152-fold. As noted above, however, with serial dilution we may expect error to compound systematically with each dilution, and indeed the value sequences in individual data sets do tend to show curves indicative of systematic pipetting error. When the model is extended to include systematic pipetting error (see Materials and Methods subsection on systematic pipetting error model), the results improve markedly (black line in Figure 3(b)), to 82.4% of valid replicates within 1.1-fold, 95.5% within 1.2-fold, and an overall geometric standard deviation improved to 1.090-fold. Fluorescein dilution provides nearly identical results: with a perfect dilution model (magenta line in Figure 3(b)), having 71.1% of valid replicates within 1.1-fold, 88.2% within 1.2-fold, and an overall geometric standard deviation of 1.148-fold, and systematic pipetting error improving the model (green line in Figure 3(b)), to 88.1% of valid replicates within 1.1-fold, 98.0% within 1.2-fold, and an overall geometric standard deviation of 1.085-fold.

Based on an analysis of the statistical properties of calibration data, we may thus conclude that the microsphere and fluorescein dilution protocols are highly robust, producing results that are precise, likely to be accurate, and readily assessed for execution quality on the basis of calibration model residuals. The LUDOX/water protocol is also highly precise and may be accurate, but its execution quality cannot be directly assessed due to its lack of residuals. The CFU protocol, on the other hand, appears likely to be highly problematic, producing unreliable and likely inaccurate calibrations.

### Reproducibility and accuracy of cell-count estimates

Reproducibility and accuracy of the calibration protocols can be evaluated through their application to calibration of fluorescence from *E. coli*, as normalized by calibrated OD measurements. Figure 4 shows the fluorescence values computed for each of the three fluorescence/OD calibration combinations, as well as for calibrated flow cytometry, excluding data with poor calibration or outlier values for colony growth or positive control fluorescence (for details see Materials and Methods on determining validity of `E. coli` data). Overall, the lab-to-lab variation was workably small, with the geometric mean of the geometric standard deviations for each test device being 2.4-fold for CFU calibration, 2.21-fold for LUDOX/water calibration, and 2.21-fold for microsphere dilution calibration. These values are quite similar to those previously reported in [9], which reported a 2.1-fold geometric standard deviation for LUDOX/water.

**Fig 4.**
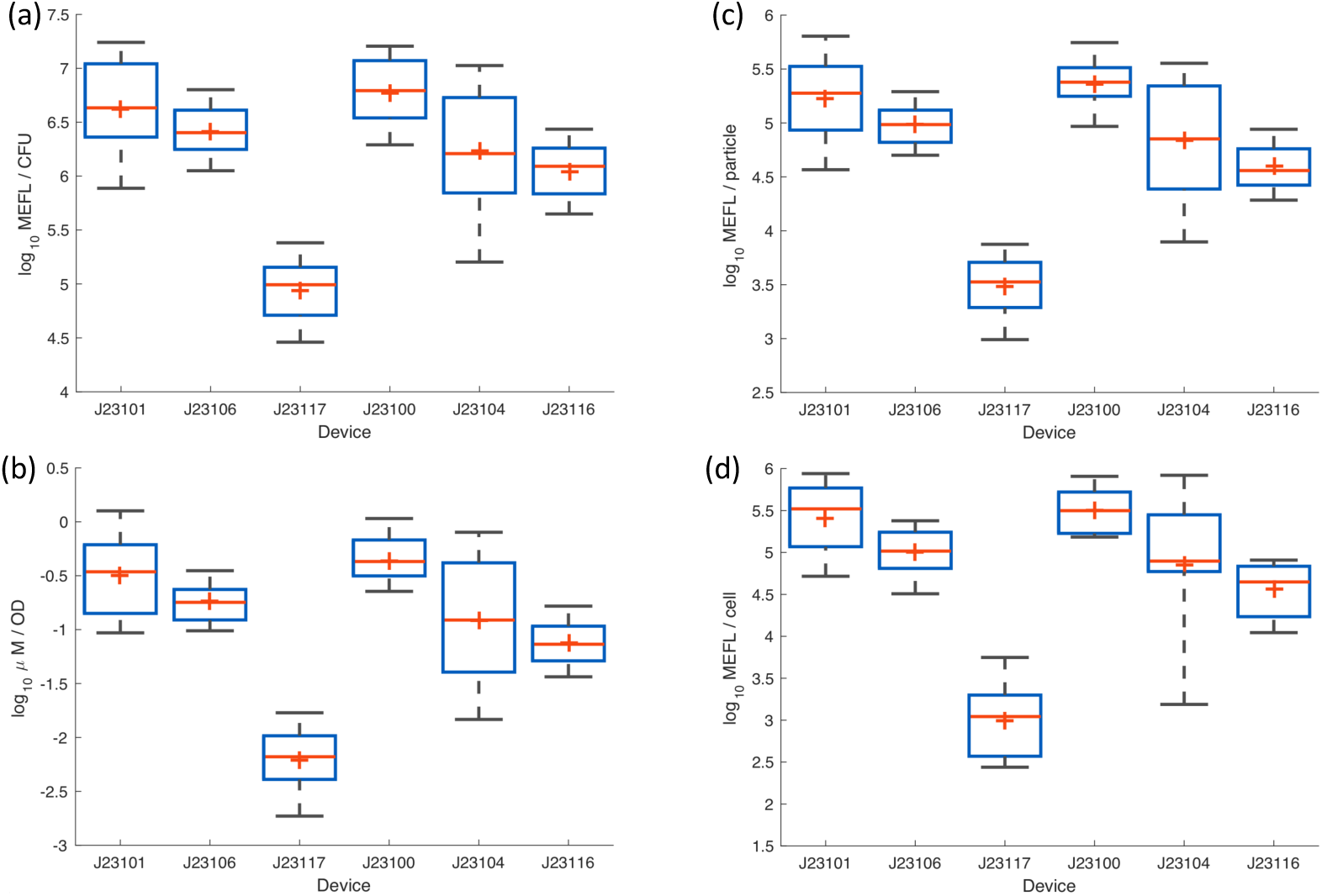
Measured fluorescence of test devices after 6 hours of growth using (a) CFU calibration, (b) LUDOX/water calibration, (c) microsphere dilution calibration, and (d) flow cytometry. In each box, red plus indicates geometric mean, red line indicates median, top and bottom edges indicate 25th and 75th percentiles, and whiskers extend from 9%–91%.

Note that these standard deviations are also dominated by the high variability observed in the constructs with J23101 and J23104, both of which appear to have suffered significant difficulties in culturing (see Supplementary Figure 2 *E. coli* Colony Growth). Omitting the problematic constructs finds variations of 2.02-fold for CFU calibration, 1.84-fold for LUDOX/water calibration, and 1.83-fold for microsphere dilution calibration. Flow cytometry in this case is also similar, though somewhat higher variability in this case, at 2.31-fold (possibly due to the much smaller number of replicates and additional opportunities for variation in protocol execution). All together, these values indicate that, when filtered using quality control based on the replicate precision and residual statistics established above, all three OD calibration methods are capable of producing highly reproducible measurements across laboratories.

To determine the accuracy of cell count estimates, we compared normalized bulk measurements (total fluorescence divided by estimated cell count) against single cell measurements of fluorescence from calibrated flow cytometry (see Materials and Methods on flow cytometry data processing for analytical details). In making this comparison, there are some differences that must be considered between the two modalities. Gene expression typically has a log-normal distribution [13], meaning that bulk measurements will be distorted upward compared to the geometric mean of log-normal distribution observed with the single-cell measurements of a flow cytometer. In this experiment, for typical levels of cell-to-cell variation observed in *E. coli*, this effect should cause the estimate of per-cell fluorescence to be approximately 1.3-fold higher from a plate reader than a flow cytometer. At the same time, non-cell particles in the culture will tend to distort fluorescence per cell estimates in the opposite direction for bulk measurement, as these typically contribute to OD but not fluorescence in a plate reader, but are gated out of flow cytometry data. With generally healthy cells in log-phase growth, however, the levels of debris in this experiment are expected to be relatively low. Thus, these two differences are likely to both be small and in opposite directions, such that we should still expect the per-cell fluorescence estimates of plate reader and flow cytometry data to closely match if accurately calibrated.

Of the three OD calibration methods, the LUDOX/water measurement is immediately disqualified as it calibrates only to a relative OD, and thus cannot produce comparable units. Comparison of CFU and microsphere dilution to flow cytometry is shown in Figure 5. The CFU-calibrated measurements are far higher than the values produced by flow cytometry, a geometric mean of 28.4-fold higher, indicating that this calibration method badly underestimates the number of cells. It is unclear the degree to which this is due to known issues of CFU, such as cells adhering into clumps, as opposed to the problems with imprecision noted above or yet other possible unidentified causes. Whatever the cause, however, CFU calibration is clearly problematic for obtaining anything like an accurate estimate of cell count.

**Fig 5.**
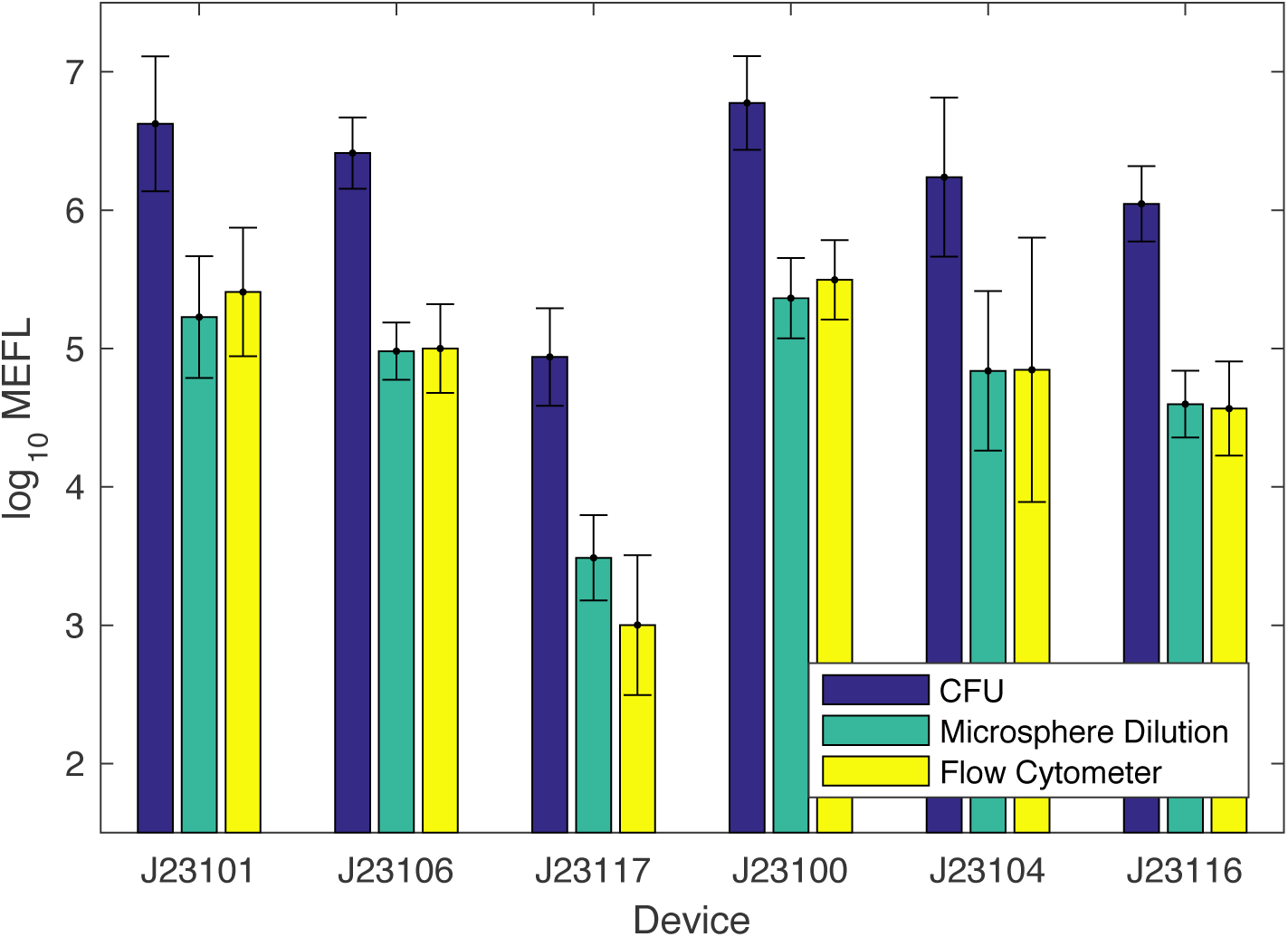
Fluorescence per cell after 6 hours of growth, comparing calibrated flow cytometry to estimates using cell count from CFU and microsphere dilution protocols (LUDOX/water is not shown as the units it produces are not comparable). Microsphere dilution produces values extremely close to the ground truth provided by calibrated flow cytometry, whereas the CFU protocol produces values more than an order of magnitude different, suggesting that CFU calibration greatly underestimates the number of cells in the sample. Bars show geometric mean and standard deviation.

**Fig 6.**
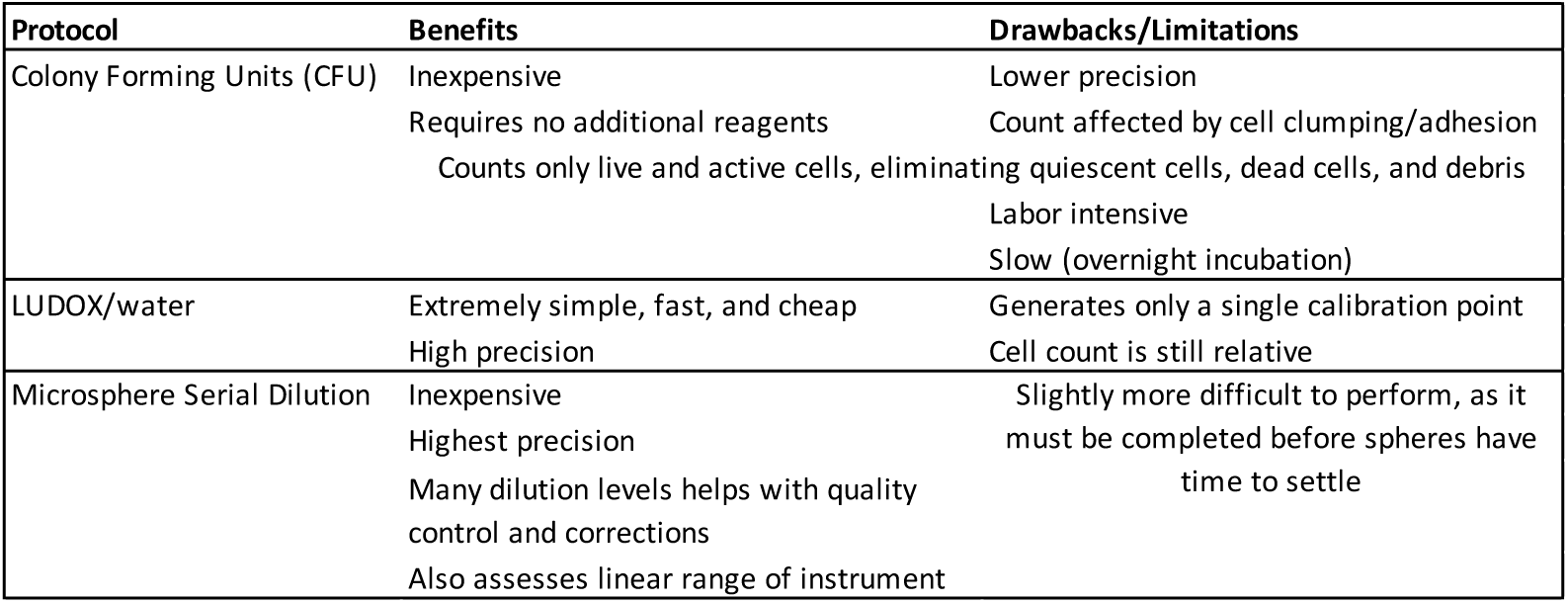
Summary of the benefits and drawbacks of the three calibration protocols.

Microsphere dilution, on the other hand, produces values that are remarkably close to those for flow cytometry, a geometric mean of only 1.07-fold higher, indicating that this calibration method is quite accurate in estimating cell count. Moreover, we may note that the only large difference between values comes with the extremely low fluorescence of the J23117 construct, which is unsurprising given that flow cytometers generally have a higher dynamic range than plate readers, allowing better sensitivity to low signals.

## Discussion

Reliably determining the number of cells in a liquid culture has remained a challenge in biology for decades. For the field of synthetic biology, which seeks to engineer based on standardized biological measurements, it was critical to find a solution to this challenge. Here, we have compared the most common method for calibrating OD to cell number (calculation of CFU) to two alternative methods of calibration: LUDOX/water and microsphere serial dilution. The qualitative and quantitative benefits and drawbacks of these three methods for OD calibration are summarized in Table 6.

These three protocols are all inexpensive, with the reagent cost for both LUDOX/water and microsphere serial dilution being less than $0.10 US. The CFU protocol has well-known issues of cell clumping and slow, labor-intensive execution, and counts only live and active cells, which can be either a benefit or a limitation depending on circumstances. Additionally, the CFU counts in this study exhibited a remarkably high level of variability, which may call into question the use of the CFU method as the a standard for determining cell counts. This observed variability is not without precedent—prior work has also demonstrated *E. coli* CFU counting performing poorly on measures of reproducibility and repeatability in an interlaboratory study [14].

The microsphere protocol, on the other hand, has no major drawbacks and provides a number of significant benefits. First, the microsphere protocol is highly robust and reliable, particularly compared to CFU assays. Second, failures are much easier to diagnose with the microsphere protocol, since it has many distinct levels that can be compared. This is particularly significant when compared to the LUDOX/water protocol, which only provides a single calibration point at low absorbance (and thus susceptible to instrument range issues), and to the CFU protocol, where failures may be difficult to distinguish from inherent high variability. With the microsphere protocol, on the other hand, some failures such as systematic dilution error and instrument saturation can not only be detected, but also modeled and corrected for. Finally, the microsphere protocol also permits a unit match between plate reader and flow cytometry measurements (both in cell number and in fluorescence per cell), which is highly desirable, allowing previously impossible data fusion between these two complementary platforms (e.g., to connect high-resolution time-series data from a plate reader with high-detail data about population structure from a flow cytometer). Accordingly, based on the results of this study, we recommend the adoption of silica microsphere calibration for robust estimation of bacterial cell count.

With regards to future opportunities for extension, we note that these methods seem likely to be applicable to other instruments that measure absorbance (e.g., spectrophotometers, automated culture flasks) by appropriately scaling volumes and particle densities. Similarly it should be possible to adapt to other cell types by selecting other microspheres with appropriately adjusted diameters and materials for their optical properties, and a wide range of potential options are already readily available from commercial suppliers. Finally, further investigation would be valuable for more precisely establishing the relationship between cell count and particle count. It would also be useful to quantify the degree to which the estimates are affected by factors such as changing optical properties associated with cell state, distribution, shape, and clustering, and to investigate means of detecting and compensating for such effects.

## Materials and Methods

Participating iGEM teams measured OD and fluorescence among the same set of plasmid-based devices, according to standardized protocols. In brief, teams were provided a test kit containing the necessary calibration reagents, a set of standardized protocols, and pre-formatted Excel data sheets for data reporting. Teams provided their own plate reader instruments, consumables/plasticware, competent *E. coli cells*, PBS, water, and culture medium. First, teams were asked to complete a series of calibration measurements by measuring LUDOX and water, and also making a standard curve of both fluorescein and silica microspheres. Next, each team transformed the plasmid devices into *E. coli* and selected transformants on chloramphenicol plates. They selected two colonies from each plate to grow as liquid cultures overnight, then the following day diluted their cultures and measured both fluorescence and OD after 0 and 6 hours of growth. Some of these cultures were also used to make serial dilutions for the CFU counting experiment. Teams were asked to report details of their instrumentation, E. coli strains used, and any variations from the protocol using an online survey.

Additional details are available in the Supplementary Information.

### Calibration Materials

The following calibration materials were provided to each team as a standard kit:

- 1 ml of LUDOX CL-X (Sigma-Aldrich)
- 1.00e-8 moles fluorescein (Sigma-Aldrich).
- 300 *µ*l of 0.961um diameter monodisperse silica beads (Cospheric) in ddH_2_0, prepared to contain 3.00e8 beads.

Fluorescein samples tubes were prepared with 1.00e-8 moles fluorescein in solution in each tube, which was then vacuum dried for shipping. Resuspension in 1 ml PBS would thus produce a solution with initial concentration of 10 *µ*M fluorescein.

Each team providing flow cytometry data also obtained their own sample of SpheroTech RCP-30-5A Rainbow Calibration Particles (SpheroTech). A sample of this material is a mixture of particles with eight levels of fluorescence, which should appear as up to eight peaks (typically some are lost to saturation on the instrument). Teams used various different lots, reporting the lot number to allow selection of the appropriate manufacturer-supplied quantification for each peak.

### Constructs, Culturing, and Measurement Protocols

The genetic constructs supplied to each team for transformation are provided in Supplementary Data 1 DNA Constructs. The protocol for plate readers, exactly as supplied to each participating team, is provided in Supplementary Note: Plate Reader and CFU Protocol. The supplementary protocol for flow cytometry is likewise provided in Supplementary Note: Flow Cytometer Protocol.

### Criteria for Valid Calibrant Replicates

For purpose of analyzing the precision of calibrants, the following criteria were used to determine which replicate sets are sufficiently valid for inclusion of analysis:

- **CFU:** A dilution level is considered valid if at least 4 of the 12 replicate plates have a number of colonies that are greater than zero but not too numerous to count. A calibration set is considered valid if there is at least one valid dilution level. Of the 244 data sets, 241 are valid and 3 are not valid.
- **LUDOX/water:** A LUDOX/water calibration is considered valid if it fits the acceptance criteria in Supplementary Note: Data Acceptance Criteria, meaning that all 244 are valid.
- **Microsphere dilution and fluorescein dilution:** For both of these protocols, a dilution level is considered locally valid if the measured value does not appear to be either saturated high or low. High saturation is determined by lack of sufficient slope from the prior level, here set to be at least 1.5x, and low saturation by indistinguishability from the blank replicates, here set to be anything less than 2 blank standard deviations above the mean blank. The valid range of dilution levels is then taken to be the longest continuous sequence of locally valid dilution levels, and the calibration set considered valid overall if this range has at least 3 valid dilution levels.
  – For microsphere dilution, of the 244 data sets, 235 are valid and 9 are not valid—one due to being entirely low saturated, the others having inconsistent slopes indicative of pipetting problems. Supplementary Figure 1 Length of Valid Sequence(a) shows that most microsphere dilution data sets have the majority of dilution levels valid, but that only about one tenth are without saturation issues.
  – For fluorescein dilution, of the 244 data sets, 243 are valid and 1 is not valid, having an inconsistent slope indicative of pipetting problems. Supplementary Figure 1 Length of Valid Sequence(a) shows that the vast majority of fluorescein dilution data sets are without any saturation issues.

### Unit Scaling Factor Computation

#### CFU

The scaling factor *S*_*c*_ relating CFU/ML to Abs600 is computed as follows:

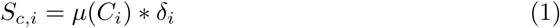

where *µ*(*C*_*i*_) is the mean number of colonies for dilution level *i* and *δ*_*i*_ is the dilution fold for level *i*. For the specific protocol used, there are three effective dilution factors, 1.6*e*5, 1.6*e*6, and 1.6*e*7 (including a 2-fold conversion between 200*µl* and 100*µl* volumes).

The overall scaling factor *S*_*c*_ for each data set is then taken to be:

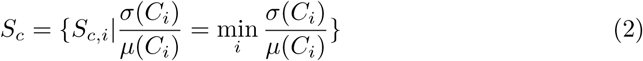

i.e., the scaling factor for the valid level with the lowest coefficient of variation.

The residuals for this fit are then *S*_*c,i*_*/S*_*c*_ for all other valid levels.

#### LUDOX/Water

The scaling factor *S*_*l*_ relating standard OD to Abs600 is computed as follow:

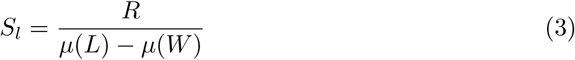

where *R* is the measured reference OD in a standard cuvette (in this case 0.063 for LUDOX CL-X), *µ*(*L*) is the mean Abs600 for LUDOX CL-X samples and *µ*(*W*) is the mean Abs600 for water samples.

No residuals can be computed for this fit, because there are two measurements and two degrees of freedom.

#### Microsphere Dilution and Fluorescein Dilution

The scaling factors *S*_*m*_ relating microsphere count to Abs600 and *S*_*f*_ for relating molecules of fluorescein to arbitrary fluorescent units are both computed in the same way. These are transformed into scaling factors in two ways, either as the mean conversion factor *S*_*µ*_ or as one parameter of a fit to a model of systematic pipetting error *S*_*p*_.

##### Mean Conversion Factor

If we ignore pipetting error, then the model for serial dilution has an initial population of calibrant *p*_0_ that is diluted *n* times by a factor of *α* at each dilution, such that the expected population of calibrant for the *i*th dilution level is:

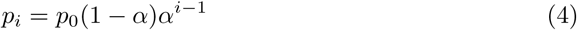

In the case of the specific protocols used here, *α* = 0.5. For the microsphere dilution protocol used, *p*_0_ = 3.00*e*8 microspheres, while for the fluorescein dilution protocol used, *p*_0_ = 6.02*e*14 molecules of fluorescein.

The local conversion factor *S*_*i*_ for the *i*th dilution is then:

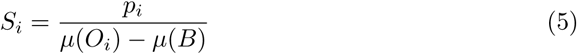

where *µ*(*O*_*i*_) is the mean of the observed values for the *i*th dilution level and *µ*(*B*) is the mean observed value for the blanks.

The mean conversion factor is thus:

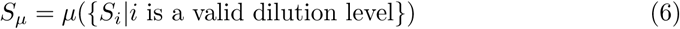

i.e., the mean over local conversion factors for valid dilution levels.

The residuals for this fit are then *S*_*i*_*/S*_*µ*_ for all valid levels.

##### Systematic Pipetting Error Model

The model for systematic pipetting error modifies the intended dilution factor *α* with the addition of an unknown bias *β*, such that the expected biased population *b*_*i*_ for the *i*th dilution level is:

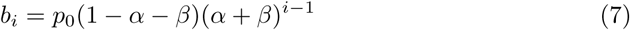

We then simultaneously fit *β* and the scaling factor *S*_*p*_ to minimize the sum squared error over all valid dilution levels:

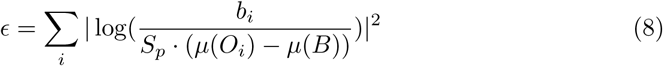

where *ϵ* is sum squared error of the fit and *x*_*n*_ is the mean corrected arbitrary unit value of the *n*th titration stage.

The residuals for this fit are then the absolute ratio of fit-predicted to observed net mean 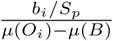 for all valid levels.

#### Application to *E. coli* data

The Abs600 and fluorescence a.u. data from *E. coli* samples are converted into calibrated units by subtracting the mean blank media values for Abs600 and fluorescence a.u., then multiplying by the corresponding scaling factors for fluorescein and Abs600.

### Criteria for Valid *E. coli* Data

For analysis of *E. coli* culture measurements, a data set was only eligible to be included if both its fluorescence calibration and selected OD calibration were above a certain quality threshold. The particular values used for the four calibration protocols were:

- **CFU:** Coefficient of variation for best dilution level is less than 0.5.
- **LUDOX/water:** Coefficient of variation for both LUDOX and water are less than 0.1.
- **Microsphere dilution:** Systematic pipetting error has geometric mean absolute residual less than 1.1-fold.
- **Fluorescein dilution:** Systematic pipetting error has geometric mean absolute residual less than 1.1-fold.

Measurements of the cellular controls were further used to exclude data sets with apparent problems in their protocol: those with a mean positive control value more than 3-fold different than the median mean positive control.

Finally, individual samples without significant growth were removed, that being defined as all that are either less than the 25% of the 75th percentile Abs600 measurement in the sample set or less than 2 media blank standard deviations above the mean media blank in the sample set.

### Flow Cytometry Data Processing

Flow cytometry data was processed using the TASBE Flow Analytics software package [15]. A unit conversion model from arbitrary units to MEFL was constructed per the recommended best practices of TASBE Flow Analytics for each data set using the bead sample and lot information provided by each team:

- Gating was automatically determined using a two-dimensional Gaussian fit on the forward-scatter area and side-scatter area channels for the first negative control (Supplementary Figure 3 Example of Flow Cytometry Gating).
- The same negative control was used to determine autofluorescence for background subtraction.
- As only a single green fluorescent protein was used, there was no need for spectral compensation or color translation.

This color model was then applied to each sample to filter events and convert GFP measurements from arbitrary units to MEFL, and geometric mean and standard deviation computed for the filtered collection of events.

## Supporting information

Supplementary Note: iGEM Interlab Study Contributors

Supplementary Note: Plate Reader and CFU Protocol

Supplementary Note: Flow Cytometer Protocol

Supplementary Note: Data Acceptance Criteria

Supplementary Data 1 DNA Constructs

Supplementary Data 2 Complete Data

Supplementary Figure 1 Length of Valid Sequence

Supplementary Figure 2 E. coli Colony Growth

Supplementary Figure 3 Example of Flow Cytometry Gating

## Supplementary Information

**Supplementary Note: iGEM Interlab Study Contributors**

**List of all authors in the iGEM Interlab Study Consortium**

**Supplementary Note: Plate Reader and CFU Protocol**

**Protocol specification provided for collecting plate reader and CFU data in the 2018 iGEM Interlab Study.**

**Supplementary Note: Flow Cytometer Protocol**

**Addendum protocol provided for flow cytometry data collection.**

**Supplementary Note: Data Acceptance Criteria**

**Quality control criteria for acceptance of data in 2018 iGEM Interlab Study.**

**Supplementary Data 1 DNA Constructs**

**File containing DNA constructs for the 2018 iGEM Interlab Study.**

**Supplementary Data 2 Complete Data**

**JSON files containing of all input data sets, plus all results of analysis per Materials and Methods above. Team names are omitted in order to anonymize data sets. For flow cytometry data, only the per-sample statistical summary of each sample is included.**

**Supplementary Figure 1 Length of Valid Sequence**

**Distribution of lengths of valid sequence of dilution levels for microspheres (a) and fluorescein (b).**

**Supplementary Figure 2** *E. coli* **Colony Growth**

**Fraction of well-grown colonies for each test construct in** *E. coli***: the constructs incorporating J23101 and J23104 presented major problems in culturing for many teams, as did the J23100 construct to a lesser degree.**

**Supplementary Figure 3 Example of Flow Cytometry Gating**

**Example of Gaussian mixture model determination of gating from negative control:**

## Acknowledgments

Partial support for this work was provided by NSF Expeditions in Computing Program Award #1522074 as part of the Living Computing Project.

This document does not contain technology or technical data controlled under either the U.S. International Traffic in Arms Regulations or the U.S. Export Administration Regulations.

## Author Contributions

- Conceptualization: J.B., N.G.F., T.H-A., V.S.-1, G.S.B., R.B-T., M.G., D.K., J.M., C.T.W.
- Data curation: J.B., N.G.F., T.H-A., V.S.-1
- Formal analysis: J.B.
- Investigation: Experimental data gathered by iGEM Interlab Study Contributors (all authors listed in Supplementary Note: iGEM Interlab Study Contributors)
- Methodology: J.B., N.G.F., T.H-A., V.S.-1, G.S.B., R.B-T., M.G., D.K., J.M., V.S.-2, A.S., C.T.W.
- Project administration: J.B., N.G.F., T.H-A.
- Resources: T.H-A., V.S.-1, A.S.
- Software: J.B.
- Writing (original draft): J.B., N.G.F.
- Writing (review & editing): J.B., N.G.F., T.H-A., G.S.B., J.M., C.T.W., V.S.-2,

## Competing Interests

The authors declare no competing interests.

