## Supplementary Note: iGEM Interlab Study Contributors for "Robust Estimation of Bacterial Cell Count from Optical Density"

Consortium authors include all persons identified by contributing teams as deserving co-authorship credit. Contributors are listed alphabetically within team, and teams alphabetically.

Team names are given as identified in iGEM records: full details of each team's institution and additional members may be found online in the iGEM Foundation archives at:

`http://2018.igem.org/Team:`*name*

e.g.: full information on the ETH_Zurich team may be found at:

`http://2018.igem.org/Team:ETH_Zurich`

The iGEM 2018 Interlab Contributors comprise a total of 1394 authors from 244 institutions.

- **Aachen:** Meryem Pehlivan, Biel Badia Roige
- **Aalto-Helsinki:** Tiu Aarnio, Samu Kivisto, Jessica Koski, Leevi Lehtonen, Denise Pezzutto, Pauliina Rautanen
- **AHUT_China:** Weixin Bian, Zhiyuan Hu, Zhihao Liu, Zi Liu, Liang Ma, Luyao Pan, Zichen Qin, Huichao Wang, Xiangxuan Wang, Hao Xu, Xia Xu
- **Aix-Marseille:** Yorgo El Moubayed
- **ASTWS-China:** Shan Dong, Choco Fang, Hanker He, Henry He, Fangliang Huang, Ruyi Shi, Cassie Tang, Christian Tang, Shirly Xu, Calvin Yan
- **Athens:** Natalia Bartzoka, Eleni Kanata, Maria Kapsokefalou, Xanthi-Leda Katopodi, Eleni Kostadima, Ioannis V. Kostopoulos, Stylianos Kotzastratis, Antonios E. Koutelidakis, Vasilios Krokos, Maria Litsa, Ioannis Ntekas, Panagiotis Spatharas, Ourania E. Tsitsilonis, Anastasia Zerva
- **Austin_LASA:** Vidhya Annem, Eli Cone, Noel Elias, Shreya Gupta, Kendrick Lam, Anna Tutuianu
- **Austin_UTexas:** Dennis M. Mishler, Bibiana Toro
- **Baltimore_BioCrew:** Akinwumi Akinfenwa, Frank Burns, Heydy Herbert, Melissa Jones, Sarah Laun, Shikei Morrison, Zion Smith
- **BCU:** Zhao Peng, Zhou Ziwei
- **BFSUICC-China:** Rui Deng, Yilin Huang, Tingyue Li, Yingqi Ma, Zhiyuan Shen, Chenxi Wang, Yuyao Wang, Tianyan Zhao
- **BGIC-Global:** Yusen Lang, Yuteng Liang, Xueyao Wang, Yi Wu
- **BGU_Israel:** Dror Aizik, Sagi Angel, Einan Farhi, Nitzan Keidar, Eden Oser, Mor Pasi
- **Bielefeld-CeBiTec:** Jorn Kalinowski, Matthias Otto, Johannes Ruhnau

- **Bilkent-UNAMBG:** Hande Cubukcu, Mehmet Ali Hoskan, Ilayda Senyuz
- **BioIQS-Barcelona:** Jordi Chi, Antoni Planas Sauter, Magda Faijes Simona
- **BioMarvel:** Sumin Byun, Sungwoo Cho, Goeun Kim, Yeonjae Lee, Sangwu Lim, Hanyeol Yang
- **BIT:** Tian Xin, Zhang Yaxi, Peng Zhao
- **BIT-China:** Weitang Han, Fa He, Yuna He, Nuonan Li, Xiaofan Luo
- **BJRS_China:** Cheng Boxuan, Hu Jiaqi, Yang Liangjian, Li Wanji, Chen Xinguang, Liu Xinyu
- **BNDS_CHINA:** Zishi Wu, Yukun Xi, Xilin Yang, Yuchen Yang, Zhuoyi Yang, Yihao Zhang, Yuezhang Zhou
- **BNU-China:** Yue Peng, Liu Yadi, Shaobo Yang, Jiang Yuanxu, Kecheng Zhang
- **BOKU-Vienna:** Doris Abraham, Theresa Heger
- **BostonU:** Cass Leach, Kevin Lorch, Linda Luo
- **British_Columbia:** Alex Gaudi, Anthony Ho, Morris Huang, Christine Kim, Luxcia Kugathasan, Kevin Lam, Catherine Pan, Ariel Qi, Cathy Yan
- **Calgary:** Kaitlin Schaaf, Cassandra Sillner
- **Cardiff_Wales:** Ryan Coates, Hannah Elliott, Emily Heath, Evie McShane, Geraint Parry, Ali Tariq, Sophie Thomas
- **CCU_Taiwan:** Ching-Wei Chen, Yu-Hong Cheng, Chia-Wei Hsu, Chin-Hsuan Liao, Wei-Ting Liu, Yu-Cheng Tang, Yu-Hsin Tang, Zon En Yang
- **CDHSU-CHINA:** Liu Jian, Caidian Li, Chenyi Lin, Guozheng Ran, Zhouyan Run, Weiyu Ting, Zhangxiang Yong, Liuhong Yu
- **Chalmers-Gothenburg:** Andrea Clausen Lind, Axel Norberg, Amanda Olmin, Jacob Sjolin, Agnes Torell, Cecilia Trivellin, Francisco Zorrilla, Philip Gorter de Vries
- **CIEI-BJ:** Haolun Cheng, Jiarong Peng, Zhenyu Xiong
- **CMUQ:** Dina Altarawneh, Sayeda Sakina Amir, Sondoss Hassan, Annette Vincent
- **CO_Mines:** Ben Costa, Isabella Gallegos, Mitch Hale, Matt Sonnier, Kathleen Whalen
- **ColumbiaNYC:** Max Elikan, Sean Kim, Jaewon You
- **Cornell:** Rahul Rambhatla, Ashwin Viswanathan
- **CPU_CHINA:** Hong Tian, Huandi Xu, Wanli Zhang, Shuyao Zhou
- **CSU_CHINA:** Liu Jiamiao, Xiao Jiaqi
- **CSU_Fort_Collins:** Darilyn Craw, Marley Goetz, Neil Rettedal, Hayden Yarbrough
- **Delgado-Ivy-Marin:** Christopher Ahlgren, Brett Guadagnino, James Guenther, Juilanne Huynh
- **DLUT_China:** Zhien He, Huan Liu, Yuansheng Liu, Mingbo Qu, Li Song, Chao Yang, Jun Yang, Xianqi Yin, Yuanzhen Zhang, Jianan Zhou, Lihan Zi
- **DLUT_China_B:** Zhu Jinyu, Xu Kang, Peng Xilei, Han Xue, Shu Xun

- **DNHS_SanDiego:** Priyanka Babu, Arushi Dogra, Pranav Thokachichu
- **DTU-Denmark:** David Faurdal, Joen Haahr Jensen, Jacob Mejlsted, Lina Nielsen, Tenna Rasmussen
- **Duesseldorf:** Jennifer Denter, Kai Husnatter, Ylenia Longo
- **Ecuador:** Juan Carlos Luzuriaga, Eduardo Moncayo, Natalia Torres Moreira, Jennifer Tapia
- **ECUST:** Tang Dingyue, Zhao Jingjing, Xu Wenhao, Teng Xinyu, Hong Xiujing
- **Edinburgh_OG:** Jackson DeKloe
- **Edinburgh_UG:** Ben Astles, Ugne Baronaite, Inga Grazulyte
- **Emory:** Michael Hwang, Yibo Pang
- **EPFL:** Michael Andrew Crone, Reza Hosseini, Moustafa Houmani, Daniel Zadeh, Violetta Zanotti
- **ETH_Zurich:** Oliver Andreas Baltensperger, Eline Yafele Bijman, Elisa Garulli, Jan Lukas Krusemann, Adriano Martinelli, Antonio Martinez, Tobias Vornholt
- **Evry_Paris-Saclay:** Monteil Camille, Ahavi Paul
- **Exeter:** Emily Browne, Daniel Barber James Gilman, Amy Hewitt, Sophie Hodson, Ingebjorg Holmedal, Fiona Kennedy, Juliana Sackey
- **FAU_Erlangen:** Selina Beck, Franziska Eidloth, Markus Imgold, Anna Matheis, Tanja Meerbrei, David Ruscher, Marco Schaeftlein
- **FJNU-China:** Zhu Hanrong
- **Fudan:** Mitchell Wan
- **Fudan-CHINA:** Leijie Dai, Kaifeng Jin, Sihan Wang, Xin Wang, Yi Wang, Yifan Wang, Chenhai Wu, Zixuan Zhang, Yineng Zhou
- **GDSYZX:** Liu Xinyu, Zeng Zirong
- **Georgia_State:** Rehmat Babar, Mathew Brewer, Christina Clodomir, Laura Das Neves, Amanda Iwuogo, Ari Jones, Cara Jones, Julia Kelly, Gloria Kim, Jessica Siemer, Yash Yadav
- **Gifu:** Yuichiro Ikagawa, Tatsuki Isogai, Ryo Niwa
- **GO_Paris-Saclay:** Celine Aubry, William Briand, Annick Jacq, Sylvie Lautru, Britany Marta, Clemence Maupu, Xavier Ollessa-Daragon, Kenn Papadopoulo, Mahnaz Sabeta Azad
- **GreatBay_China:** Wei Kuangyi, Yao Xiu, Chenghao Yang
- **Groningen:** Aditya Iyer, Rianne Prins, Phillip Yesley
- **GZHS-United:** Fang Lichi, Chen Zi Xuan
- **HAFS:** Kyuhee Jo, Mikyung Park, Seunghyun Park, Hojun Yoo
- **Hamburg:** Nele Burckhardt, Lea Daniels, Bjarne Klopprogge, Dustin Kruger, Oda-Emilia Meyfarth, Lisa Putthoff, Dominika Wawrzyniak
- **HBUT-China:** Xinyi Hu, Yunyi Wang
- **HebrewU:** Lior Badash, Amichai Baichman-Kass, Alon Barshap, Yonatan Friedman, Eliya Milshtein, Omri Vardi

- **HFLS_ZhejiangUnited:** Shan Dong, Yining Gu, Yuanzhe Pei, Ruyi Shi, Fan Yang, Jinshu Yang, Xueqian Zhu
- **HK_HCY_LFC:** Lam Kai Ching, Law Hiu Ching, Ng Tsz Chun, Yu Man Hin, Lai Tsz Hong, Chan Wing Lam, Yiu Choi Lam, Cheah Matthew, Cheng Tsz Ngo, Yun Shuan, Chan Tsey Wan, Tsui Shing Yan, Chong Yuk Yee, Tam Chi Yu, Yuen Wai Yu
- **HKJS_S:** Chung Tsun Ho Anson, Lee Sze Choi, Cheung Man Chun, Chan Lok Hin, Wong Chung Hin, Ng Sze Ho, Leung Chung Yin Jay, Lai Man Wai Katherine, Wong Carol Kin-ning, Lee Hong Kiu, Cheng Chak Kong, Leung Chung Wai, Yeung Wing Yan, Wong Tsz Yeung, Lee Ka Yin
- **Hong_Kong_HKU:** Tsui Shing Yan Grace, Lam Kai Ching Joe, Ng Tsz Chun Kenneth, Cheah Matthew Yun Shuan
- **Hong_Kong_HKUST:** Ferdinan Aldo, Chung Him Pang, Kam Pang So, Hei Man Wong
- **Hong_Kong_JSS:** Lai Tsz Ching, Luk Hau Ching, Ip Ning Fung, Yam Shing Fung, Lee Chi Hong, Hsiu Ou Ning, Jonathan Cheng Hon Sang
- **Hong_Kong-CUHK:** Yeung Hoi Lam Elsa, Chan Yick Hei, Lo Ho Sing, Choi Seong Wang
- **HUBU-Wuhan:** Yiheng Gu, Ziyue Rong, Haoyue Song, Pengying Wang, Yuefei Wang
- **HUST-China:** Yan Chen, Hao Qiu, Haotian Ren, Ziyang Xiao
- **HZAU-China:** Heng Heng, Xichen Rao, Ruonan Tian
- **ICT-Mumbai:** Shalini S. Deb, Yash Laxman Kamble, Ninad Kumbhojkar, Bhargav Patel, Supriya Prakash, Shamlan M.S. Reshamwala, Poorva Taskar
- **IISc-Bangalore:** Gokul, Adwaith B Uday
- **IISER-Bhopal-India:** Anubhav Basu, Rishi Gandhi, Jatin Khaimani, Arundhati Khenwar, Sandeep Raut, Tejas Somvanshi
- **IISER-Kolkata:** Diptatanu Das, Souvik Ghosh, Hrishika Rai
- **IISER-Mohali:** Nithishwer Mouroug Anand, Ashwin Kumar Jainarayanan, Pranshu Kalson, Devang Haresh Liya, Vibhu Mishra, Sveekruth Sheshagiri Pai, Madhav Pitaliya, Yash Rana, Ravineet Yadav
- **IIT_Delhi:** Neha Arora, Vasu Arora, Shubham Jain, Abhilash Patel, Saksham Sharma, Priyanka Singh
- **IIT_Kanpur:** Anushya Goenka, Rishabh Jain, Aryaman Jha, Adarsh Kumar, Abhinav Soni
- **IIT-Madras:** Sathvik Ananthakrishnan, Velvizhi Devi, Mohammed Faidh, Guhan Jayaraman, M Sagar Kittur, Nitish R Mahapatra, Sarvesh Menon, Anantha Barathi Muthukrishnan, Kailash B P, Burhanuddin Sabuwala, Mousami Shinde, Sankalpa Venkatraghavan
- **Jiangnan_China:** Weijia Liu, Zhoudi Miao, Tian Wang, Yaling Wang, Shuyan Zhang
- **Jilin_China:** Ruochen Chai, Yubin Ge, Ali Hou, Fangqi Liu, Xutong Liu, Jiangjiao Mao, Zihao Wang, Haimeng Yu, Hetian Yuan, Yang Zhan
- **JMU_Wuerzburg:** Anna Ries, Chiara Wolfbeisz
- **KAIT_JAPAN:** Toshihiro Kanaya, Yusuke Kawasaki, Tatuya Maruo, Yuya Mori, Takehito Satoh
- **KCL_UK:** Anthony Chau, Wai Yan Chu, Anatoliy Markiv, Marcos Vega-Hazas Marti, Maria Jose Ramos Medina, Deeksha Raju, Shubhankar Sinha

- **KUAS_Korea:** Youngeun Choi, Bo Sun Ryu
- **Lambert_GA:** Gaurav Byagathvalli, Ellie Kim
- **Leiden:** Marjolein Crooijmans, Jazzy de Waard, Chiel van Amstel
- **Lethbridge:** Aubrey Demchuk, Travis Haight, Dong Ju Kim, Andrei Neda, Luc Roberts, Luke Saville, Reanna Takeyasu, David Tobin
- **Lethbridge_HS:** Mina Akbary, Rebecca Avileli, Karen He, Aroma Pageni, Luke Saville, Dewuni De Silva, Nimaya De Silva, Kristi Turton, Michelle Wu, Alice Zhang
- **Lubbock_TTU:** Benjamin Chavez, Paula Garavito, Michael Latham, Jeffrey Ptak, Darron Tharp
- **Lund:** Nurul Izzati, Martin Jonsson, Nikol Labecka, Sara Palo
- **Macquarie_Australia:** Renee Beale, Dominic Logel, Areti-Efremia Mellou, Karl Myers
- **Madrid-OLM:** Alejandro Alonso, Rodrigo Hernandez Cifuentes, Borja Sanchez Clemente, Gonzalo Saiz Gonzalo, Ivan Martin Hernandez, Laura Armero Hernandez, Francisco Javier Quero Lombardero, Domingo Marquina, Guillermo Fernandez Rodriguez, Ignacio Albert Smet
- **Manchester:** Tom Butterfield, Ed Deshmukh-Reeves, Namrata Gogineni, Sam Hemmings, Ismat Kabbara, Ieva Norvaisaite, Ryan Smith
- **Marburg:** Daniel Bauersachs, Benjamin Daniel, Rene Inckemann, Alexandra Seiffermann, Daniel Stukenberg, Carl Weile
- **McGill:** Valerian Clerc, Jacqueline Ha, Stephanie Totten
- **McMaster:** Thomas Chang, Carlene Jimenez, Dhanyasri Maddiboina
- **METU_HS_Ankara:** Beliz Leyla Acar, Evrim Elcin, Tugba Inanc, Gamze Kantas, Ceyhun Kayihan, Mert Secen, Gun Suer, Kutay Ucan, Tunc Unal
- **Michigan:** Matthew Fischer, Naveen Jasti, Thomas Stewart
- **MichiganState:** Sarah Caldwell, Jordan Lee, Jessica Schultz
- **Mingdao:** Ting-Chen Chang, Pei-Hong Chen, Yu-Hsuan Cheng, Yi-Hsuan Hsu, Chan-yu Yeh
- **Minnesota:** Zhipeng Ding, Zihao Li, Savannah Lockwood, Katherine Quinn
- **Montpellier:** Leo Carrillo, Maxime Heintze, Lea Meneu, Marie Peras, Tamara Yehouessi
- **Munich:** Keno Eilers, Elisabeth Falgenhauer, Wong Hoi Kiu, Julia Mayer, Julia Mueller, Sophie von Schoenberg, Dominic Schwarz, Brigit Tunaj
- **Nanjing-China:** Zhaoqing Hu, Yansong Huang, Yuanyuan Li
- **NAU-CHINA:** Chengzhu Fang, Jiangyuan Liu, Yiheng Liu, Yaxuan Wu, Sheng Xu, Long Yuan
- **NAWI_Graz:** Marco Edelmayer, Marlene Hiesinger, Sebastian Hofer, Birgit Krainer, Andreas Oswald, Dominik Strasser, Andreas Zimmermann
- **NCHU_Taichung:** Yi-Cian Chen
- **NCTU_Formosa:** Yuan-Yao Chan, Yu-Ci Chang, Nian Ruei Deng, Chi-Yao Ku, Meng-Zhan Lee
- **NEU_China_A:** Hailong Li, Zhaoyu Liu, Guowei Song, Yuening Xiang, Hongfa Yan

- **NEU_China_B:** He Huanying, Jiang Qiaochu, Jiang Shengjuan, Peng Yujie
- **Newcastle:** Matt Burridge, Kyle Standforth, Sam Went
- **NJU-China:** Liang Chenxi, Wang Han, Zhang Qipeng, Li Yifan, Quan Yiming, Pan Yutong
- **NKU_CHINA:** Senhao Kou, Lin Luan
- **Northwestern:** Umut Akova, Liza Fitzgerald, Bon Ikwuagwu, Michael Johnson, Jacob Kurian, Christian Throsberg
- **Nottingham:** Lucy Allen, Christopher Humphreys, Daniel Partridge, Michaella Whittle, Nemira Zilinskaite
- **NPU-China:** Meixuan Lee, Weifeng Lin, Yuan Ma, Kai Wang
- **NTHU_Formosa:** Hsuan Cheng, Shumei Chi, Yi-Chien Chuang, Ray Huang, LiangYu Ko, Yu-Chun Lin
- **NTHU_Taiwan:** You-Yang Tsai, Cheng-Chieh Wang, Kai-Chiang Yu
- **NTNU_Trondheim:** Hanna Nedreberg Burud, Carmen Chen, Anne Kristin Haralsvik, Adrian Marinovic, Hege Hetland Pedersen, Amanda Sande, Vanessa Solvang
- **NTU-Singapore:** Shaw Kar Ming, Albert Praditya
- **NU_Kazakhstan:** Aiganym Abduraimova, Ayagoz Meirkhanova, Assel Mukhanova, Tomiris Mulikova
- **NUDT_CHINA:** Yanchen Gou, Chenyu Lu, Jiaxin Ma, Chushu Zhu
- **NUS_Singapore-A:** Leow Chung Yong Aaron, Tvarita Shivakumar Iyer, Wu Jiacheng, Yan Ping Lim, Beatrix Tung Xue Lin, Aaron Ramzeen, Nur Liyana Binte Ayub Ow Yong
- **NUS_Singapore-Sci:** Yah Tse Sabrina Chua, Yuhui Deborah Fong, Menglan He, Li Yang Tan
- **NWU-China:** Zhang Jiahe, Li Mingge, Li Nianlong, Li Yueyi, Cheng Yuhan
- **NYMU-Taipei:** Annabel Chang, Chih-Chiang Chen, Ryan Chou, Jude Clapper, Evelyn Lai, Yasmin Lin, Kelsey Wang, Jake Yang
- **NYU_Abu_Dhabi:** Mariam Anwar, Ibrahim Chehade, Imtiyaz Hariyani, Sion Hau, Ashley Isaac, Laura Karpauskaite, Mazin Magzoub, Daniel Obaji, Yong Rafael Song, Yejie Yun
- **OUC-China:** Kai Sun, Yunqian Zhang
- **Oxford:** Eleanor Beard, Laurel Constanti Crosby, Nicolas Delalez, Arman Karshenas, Adrian Kozhevnikov, Jhanna Kryukova, Karandip Saini, Jon Stocks, Bhuvana Sudarshan, Max Taylor, George Wadhams, Joe Windo
- **Paris_Bettencourt:** Annissa Ameziane, Darshak Bhatt, Alexis Casas, Antoine Levrier, Ana Santos, Nympha Elisa M. Sia, Edwin Wintermute
- **Pasteur_Paris:** Alice Dejoux, Deshmukh Gopaul, Lea Guerassimoff, Samuel Jaoui, Manon Madelenat, Serena Petracchini
- **Peking:** Fu Cai, Yang Jianzhao, Shi Shuyu, Li Tairan, Li Xin, Lin Yongjie, Huang Zhecheng
- **Pittsburgh:** Evan Becker, Matthew Greenwald, Vivian Hu, Tucker Pavelek, Elizabeth Pinto, Zemeng Wei

- **Purdue:** Zachary Burgland, Janice Chan, Julianne Dejoie, Kevin Fitzgerald, Zach Hartley, Moiz Rasheed, Makayla Schacht
- **Queens_Canada:** Maddison Gahagan, Ellis Kelly, Elisha Krauss
- **RDFZ-China:** Yuze Cao, Yishen Shen, Xuan Wang, Hanning Xu, Jianxiang Zhang
- **REC-CHENNAI:** Priyanka Chandramouli, Amal Jude Ashwin F, Srimathi Jayaraman, Marcia Smiti Jude, Vignesh Kumar, Hema Lekshmi, Preetha R, Khadija Rashid, Deepak Kumar S, Mohan Kumar B S
- **Rheda_Bielefeld:** Leon Michael Barrat, Jil-Sophie Dissmann, Jorn Kalinowski, Matthias Otto, Johannes Ruhnau, Fynn Stuhlweissenburg, Elisa Ueding
- **RHIT:** Ariel Bohner, Brittany Clark, Emilie Deibel, Liz Klaas, Kaylee Pate, Elisa Weber
- **Rice:** Katherine Cohen, Anna Guseva, Stefanie King, Soohyun Yoon
- **Ruia-Mumbai:** Sanika Ambre, Shilpa Bhowmick, Nishtha Pange, Komal Parab, Vainav Patel, Mitali Patil, Aishwarya Rajurkar, Mayuri Rege, Maithili Sawant, Shrutika Sawant, Anjali Vaidya
- **SBS_SH_112144:** Peicheng Ji, Fang Luo, Guanghui Ma, Xin Xu, Jiacheng Yin, Yinchi Zhou, Ke Zhu
- **SCAU-China:** Yaohua Huang, Yinpin Huang, Jiadong Li, Xuecheng Li, Hao Wang, Ken Wang, Wei Wang, Xinyu Zhang, Jiahua Zou
- **SCU-China:** Minyue Bao, Han Kang, Xiaolong Liu, Yibing Tao, Ziru Wang, Fuqiang Yang, Tianyi Zhang, Yanling Zhong
- **SCUT_ChinaB:** Jiezheng Liu, Jingang Liu, Lingling Ma, Xubo Niu, Ling Qian, Li Wang, Qingyan Yan, Nannan Zhao
- **SCUT-ChinaA:** Weixuan Chen, Yuxin Zhou
- **SDU-CHINA:** Junyang Chen
- **SFLS_Shenzhen:** Junyao Hao, Zhang HuaYue, Peilin Li, Yifei Pei, Jingting Qu, Raven Wang, Xinyue Wang, Kangjie Wu, Yuxuan Wu, Meredith Xiang, Leyi Yang, Zisang Yang, Li Zhaoting
- **ShanghaiTech:** Wenhan Fu, Zonghao Li, Weiyi Tang, Kaida Zhang
- **SHSBNU_China:** Haocong Li, Xuze Shao, Chuyi Yang, Yuanhong Zeng, Yanjun Zhou
- **SHSID_China:** Shangzhi Dong, Younji Jung, Sophie Ruojia Li, Tingting Li, Jiacheng Yu
- **SHSU_China:** Shangzhi Dong, Tingting Li, Xinyi Miao, Sibo Wang
- **SIAT-SCIE:** Yiming Ding, Jiaxi Huang, Yuqi Li, Ting Sun, Qinghe Tian, Mengxuan Wu, Jinming Xing, Xin Xiong, Yining Yan, Qiu Yihang, Jige Zhang, Yi Zhou, Zhiyu Zhou
- **SJTU-BioX-Shanghai:** Zhuoyang Chen, Peixiang He, Yirui Hong, Chia-Yi Hsiao, Zhihan Liang, Zhixiang Liu, Yuncong Ran, Shiyu Sun, Ruoyu Xia
- **SKLMT-China:** Dongyang Dong, Wenxue Zhao
- **SMMU-China:** Miao Hu, Shi Hu, Wei Shi, Shulun, Han Yan, Yusheng Ye
- **SMS_Shenzhen:** Yiquan Hong, Yuyao Pan, Yiran Song, Jinhan Zhang, Yihang Zhao
- **Sorbonne_U_Paris:** Dounia Chater, Asmaa Foda, Yanyan Li, Ursula Saade, Victor Sayous

- **SSHS-Shenzhen:** Yilin Mo, Wenan Ren, Chenxu Zeng
- **SSTi-SZGD:** Yixin Cao
- **St_Andrews:** Clarissa Czekster, Izzy Dunstan, Simon Powis, Bethany Reaney, Eva Snaith, Cam Young
- **Stanford:** Eva Frankel, Eleanor Glockner, Isaac Justice
- **Stanford-Brown-RISD:** Santosh Murugan, Leo Penny
- **Stockholm:** Chrismar Garcia, Stamatina Rentouli
- **Stony_Brook:** Priya Aggarwal, Stephanie Budhan, Woody Chiang, Dominika Kwasniak, Karthik Ledalla, Matthew Lee, Natalie Lo, Matthew Mullin, Lin Yu Pan, Jennifer Rakhimov, Robert Ruzic, Manvi Shah, Lukas Velikov, Sara Vincent
- **Stuttgart:** Philip Horz, Nadine Kuebler, Jan Notheisen
- **SUIS_Shanghai:** David Doyle, Jiajun Gu, Wenyue Hu, Shuting Yang
- **SYSU-CHINA:** Tao Kehan, Gao Menghan, Mao Xiaowen
- **SYSU-Software:** Yifei Chen, Ziqi Kang, Haochen Ni
- **SZU-China:** Junyu Chen, Lindong He, Mingyue Luo, Jiaqi Tang
- **Tacoma_RAINmakers:** Kira Boyce, James Lee, Michael Martin, Judy Van Nguyen, Leon Wan
- **Tartu_TUIT:** Artur Astapenka, Turan Badalli, Irina Borovko, Nadezhda Chulkova, Anastasia Kolosova, Artemi Maljavin, Frida Matiyevskaya, Vladislav Tuzov
- **TAS_Taipei:** Catherine Chang, Ryan Chou, Jude Clapper, Tim Ho, Yi Da Hsieh, Evelyn Lai, Leona Tsai, Kelsey Wang, Justin Wu
- **Tec-Chihuahua:** Viana Isabel Perez Domnguez, Cesar Ibrahym Rodriguez Fernandez, Daniela Olono Fierro, Anna Karen Aguilar Nunez, Jose Pablo Rascon Perez, Mario Loya Rivera, Cynthia Lizeth Gonzalez Trevizo, Maria Antonia Luna Velasco
- **Tec-Monterrey:** Carlos Javier Cordero Oropeza, Adrian Federico Hernandez Mendoza, Jose Arnulfo Juarez Figueroa, Luis Mario Leal, Samantha Ayde Pena Benavides, Victor Javier Robledo Martinez, Adriana Lizeth Rubio Aguirre, Andres Benjamin Sanchez Alvarado, Margarita Sofia Calixto Solano, Nora Esther Torres Castillo, Alejandro Robles Zamora, Esteban de la Pena Thevenet
- **TecCEM:** Karla Soto Blas, Ana Laura Torres Huerta, Armando Cortes Resendiz
- **TecMonterrey_GDL:** Frida Cruz, Fernanda Diaz, Diego Espinoza, Ana Cristina Figueroa, Ana Cecilia Luque, Roberto Portillo, Carolina Senes, Diana Tamayo, Mariano Del Toro
- **Thessaloniki:** Ioannis Alexopoulos, Alexandros Dimitriou Giannopoulos, Yvoni Giannoula, Grigorios Kyrpizidis
- **Tongji_China:** Ma Xinyue, Chen Xirui, Song Zhiwei
- **Toronto:** Nina Adler, Amalia Caballero, Carla Hamady, Ahmed Ibrahim, Jasmeen Parmar, Tashi Rastogi, Jindian Yang
- **Toulouse-INSA-UPS:** Jean Delhomme, Anthony Henras, Stephanie Heux, Yves Romeo, Marion Toanen, Camille Wagner, Paul Zanoni

- **TU_Darmstadt:** Thea Lotz, Elena Nickels, Beatrix Suss, Heribert Warzecha, Jennifer Zimmermann
- **TU-Eindhoven:** Emilien Dubuc, Bruno Eijkens, Sander Keij, Simone Twisk, Mick Verhagen, Maxime van den Oetelaar
- **TUDelft:** Alexander Armstrong, Nicole Bennis, Susan Bouwmeester, Lisa Buller, Kavish Kohabir, Monique de Leeuw, Venda Mangkusaputra, Jard Mattens, Janine Nijenhuis, Timmy Paez, Lisbeth Schmidtchen, Gemma van der Voort
- **TUST_China:** Gao Ge, Xu Haoran, Li Xiaojin
- **UAlberta:** Ejouan Agena, Ethan Agena, Scott Bath, Robert Campbell, Rochelin Dalangin, Anna Kim, Dominic Sauvageau, Irene Shkolnikov
- **UC_Davis:** Daniel Graves, Jacob Lang, Jolee Nieberding-Swanberg, Achala Rao, Ares Torres, Andrew Yao
- **UC_San_Diego:** Anser Abbas, Claire Luo
- **UCAS-China:** Xu Zepeng, Zhao Ziyi
- **UChicago:** Janice Chen, Cian Colgan, Steve Dvorkin, Rachael Filzen, Varun Patel, Allison Scott, Patricia Zulueta
- **UChile_Biotec:** Joaquin Acosta, Lucas Araya, Francisco Chavez, Sebastian Farias, Delia Garrido, Andres Marcoleta, Felipe Munoz, Paula Rivas
- **UCL:** Noelle Colant, Catherine Fan, Stefanie Frank, Jacopo Gabrielli, Paola Handal, Vitor Pinheiro, Stefanie Santamaria, Shamal Withanage, Fang Xue
- **UCLouvain:** Antoine Gerard, Marine Lefevre, Fiona Milano, Nina De Sousa Oliveira, Mathieu Parmentier, Luca Rigon
- **UConn:** Elizabeth Chamiec-Case, Ryan Chen, Peter Crowley, Shannon Doyle, Sricharan Kadimi, Toni Vella
- **UCopenhagen:** Natthawut Adulyanukosol, Theodore A Dusseaux, Victor Forman, Cecilie Hansen, Selma Kofoed, Simon Louis, Magnus Ronne Lykkegaard, Davide Mancinotti, Lasse Meyer, Stephanie Michelsen, Morten Raadam, Victoria Svaerke Rasmussen, Eirikur Andri Thormar, Attila Uslu, Nattawut leelahakorn
- **UESTC-China:** Shizhi Ding, Changyu Li, Huishuang Tan, Yinsong Xu, Jianzhe Yang
- **UFlorida:** Diego Gamoneda, Nicole Kantor, Lidimarie Trujillo-Rodriguez, Matthew Turner
- **UGA:** Stephan George, Kelton McConnell, Chynna Pollitt
- **UI_Indonesia:** Ihya Fakhrurizal Amin, Muhammad Ikhsan, Valdi Ven Japranata, Andrea Laurentius, Luthfian Aby Nurachman, Muhammad Iqbal Adi Pratama
- **UiOslo_Norway:** Yvette Dirven, Lisa Frohlich, Dirk Linke, Verena Mertes, Rebekka Rekkedal Rolfsnes, Athanasios Saragliadis
- **UIOWA:** Sandra Castillo, Sathivel Chinnathambi, Craig Ellermeier, Jennifer Farrell, Jan Fassler, Ernie Fuentes, Sean Ryan, Edward Sander
- **UIUC_Illinois:** Amie Bott, Liam Healy, Pranathi Karumanchi, Alex Ruzicka, Ziyu Wang

- **ULaval:** Gabriel Byatt, Philippe C Despres, Alexandre Dube, Pascale Lemieux, Florian Lepetit, Louis-Andre Lortie, Francois D Rouleau
- **ULaVerne_Collab:** Seth Barrington, Cynthia Basulto, Sabrina Delgadillo, Karen De Leon, Micah Madrid, Catherosette Meas, Angelica Sabandal, Magaly Aguirre Sanchez, Jennifer Tsui, Noble Woodward
- **UMaryland:** Rohith Battina, Jess Boyer, Arjun Cherupalla, Jason Chiang, Mary Heng, Collin Keating, Tommy Liang, Chun Kit Loke, Jacob Premo, Keerthana Srinivasan, John Starkel, Daniel Zheng
- **UNebraska-Lincoln:** Gabe Astorino, Rachel Van Cott, Jintao Guo, Drew Kortus, Wei Niu
- **Unesp_Brazil:** Paulo J. C. Freire, Danielle Biscaro Pedrolli, Nathan Vincius Ribeiro, Bruna Fernandes Silva, Nadine Vaz Vanini, Mariana Santana da Mota, Larissa de Souza Crispim
- **UNSW_Australia:** Tyler Chapman, Tobias Gaitt, Megan Jones, Emily Watson
- **UPF_CRG_Barcelona:** Guillem Lopez-Grado, Laura Sans
- **Uppsala:** Matilda Brink, Varshni Rajagopal, Elin Ramstrom
- **US_AFRL_CarrollHS:** Anna Bete, Yazmin Camacho, Jonah Carter, Christina Davis, Jason Dong, Amy Ehrenworth, Michael Goodson, Chris Guptil, Max Herrmann, Chia Hung, Hayley Jesse, Rachel Krabacher, Dallas McDonald, Peter Menart, Travis O'Leary, Laura Polanka, Andrea Poole, Vanessa Varaljay
- **USMA-West_Point:** Alana Appel, John Cave, Liz Huuki, Matt McDonough, Channah Mills, Alex Mitropoulos, James Pruneski, Ken Wickiser
- **USP-Brazil:** Felipe Xavier Buson, Vinicius Flores, Guilherme Meira Lima, Caio Gomes Tavares Rosa
- **UST_Beijing:** Guanke Bao, Haitao Dong, Zhi Luo, Jiarong Peng
- **USTC:** Yongyan An, Cheng Cheng, Zhenyu Jiang, Linzhen Kong, Chenfei Luo, Liudong Luo, Yingying Shi, Erting Tang, Ping Wang, Yuyang Wang, Guiyang Xu, Wenfei Yu, Bonan Zhang, Qian Zhang
- **UT-Knoxville:** David Garcia, Nannan Jiang, Brandon Kristy, Ralph Laurel, Karl Leitner, Frank Loeffler, Steven Ripp, Morgan Street
- **Utrecht:** Khadija Amheine, Felix Bindt, Meine Boer, Mike Boxem, Jolijn Govers, Seino Jongkees, Lorenzo Pattiradjawane, Pim Swart, Helen Tsang, Floor de Graaf, Marjolijn ten Dam, Franca van Heijningen
- **Valencia_UPV:** Yadira Boada, Alejandro Vignoni
- **Vilnius-Lithuania:** Valentas Brasas, Aukse Gaizauskaite, Gabrielius Jakutis, Simas Jasiunas, Ieva Juskaite, Justas Ritmejeris, Dovydas Vaitkus, Tomas Venclovas, Kornelija Vitkute, Hanna Yeliseyeva, Kristina Zukauskaite, Justina Zvirblyte
- **Vilnius-Lithuania-OG:** Laurynas Karpus, Ignas Mazelis, Irmantas Rokaitis
- **Virginia:** Ngozi D. Akingbesote, Dylan Culfogienis, William Huang, Kevin Park
- **Warwick:** Janvi Ahuja, Christophe Corre, Gurpreet Dhaliwal, Rhys Evans, Kurt Hill, Olivor Holman, Alfonso Jaramillo, Alizah Khalid, Jack Lawrence, Laura Mansfield, James O'Brien, June Ong, Satya Prakash, Jonny Whiteside
- **Washington:** Karl Anderson, Emily Chun, Grace Kim, Aerilynn Nguyen, Chemay Shola, Dorsa Toghani, Angel Wong, Joanne Wong, Jay Yung

- **WashU_StLouis:** Elizabeth Johnson, Divangana Lahad, Kyle Nicholson, Havisha Pedamallu, Cam Phelan
- **Waterloo:** Clara Fikry, Leah Fulton, Nicole Lassel, Dylan Perera, Marina Robin, Nicolette Shaw
- **Westminster_UK:** Kyle Bowman, Sarah Coleman, Kristian Emilov, Camila Gaspar, Jenaagan Jenakendran, Sara Mubeen, Marko Obrvan, Caroline Smith
- **WHU-China:** Tang Bo, Du Liaoqi, Chang Tianyi, Xing Yuan, Qing Yue
- **William_and_Mary:** Stephanie Do, Xiangyi Fang, Ethan Jones, Jessica Laury, Wukun Liu, Adam Oliver, Lillian Parr, Margaret Saha, Chengwu Shen, Tinh Son, Julia Urban, Yashna Verma, Hanmi Zhou
- **Worldshaper-Wuhan:** Shan Dong, Zhengguo Hao, Yi Kuang, Ting Liu, Rui Zhou
- **WPI_Worcester:** Beck Arruda, Natalie Farny, Mei Hao, Camille Pearce, Alex Rebello, Arth Sharma, Kylie Sumner, Bailey Sweet
- **XJTLU-CHINA:** Junliang Lin
- **XJTU-China:** Du Mengtao, Fan Peiyao, Fang Xinlei
- **XMU-China:** Niangui Cai, Junhong Chen, Yousi Fu, Yunyun Hu, Ye Qiang, Qiupeng Wang, Ruofan Yang, Chen Yucheng, Jiyang Zheng
- **Yale:** Kevin Chang, Cecily Gao, Farren Isaacs, Kevin Li, Ricardo Moscoso, Jaymin Patel, Lauren Telesz, Alice Tirad
- **ZJU-China:** Qinhao Cao, Xinhua Feng, Yinjing Lu, Xianyin Zhang, Xuanhao Zhou
- **ZJUT-China:** Dongchang Sun, Zhe Yuan, Jiajie Zhou
