## Supplementary Note: Plate Reader and CFU Protocol for "Robust Estimation of Bacterial Cell Count from Optical Density"

### iGEM 2018 InterLab Study Protocol

#### Before You Begin

Read through this entire protocol sheet carefully before you start your experiment and prepare any materials you may need. In order to improve reproducibility, **we are requiring all participating teams to use plate readers to take measurements of fluorescence and absorbance**. If you do not have access to a plate reader with those capabilities, you may collaborate with another team. If the plate reader requirement is a significant barrier for your team, you can still participate in the InterLab study. Contact the iGEM Measurement Committee at measurement at igem dot org to discuss your situation.

Before beginning your experiments, it will be helpful to gather the following information about your plate reader, as you will be asked to provide this information when submitting your data to iGEM HQ:

- ❏ Instrument brand and model ______________________________________________
- ❏ Can your instrument measure both absorbance and fluorescence? _______________
- ❏ Does your instrument have pathlength correction, and if yes can it be disabled? ____________________________________________________________________
- ❏ Does your instrument have variable temperature settings, and if yes can this be set to room temperature? ______________________________________________________
- ❏ What filters does your instrument have for measuring GFP? You will need information about the bandpass width (530 nm / 30 nm bandpass, 25-30nm width is recommended), excitation (485 nm is recommended) and emission (520-530 nm is recommended) of this filter. ______________________________________________
- ❏ Does your instrument use top or bottom optics (i.e. does your plate reader read samples from the top of the plate or the bottom)? ____________________________

You will need all of the following supplies and reagents to complete this entire protocol. Please take a moment to check that you have all of these supplies and reagents before you begin:

- ❏ Measurement Kit (provided with the iGEM distribution shipment) containing:
  - ❏ 1ml LUDOX CL-X
  - ❏ 150 µL Silica Bead (microsphere suspension)
  - ❏ Fluorescein (powder, in amber tube)

- ❏ iGEM Parts Distribution Kit Plates (you will obtain the test devices from the parts kit plates)
- ❏ 1x PBS (phosphate buffered saline, pH 7.4 - 7.6)
- ❏ dd$H_2O$ (ultrapure filtered or double distilled water)
- ❏ Competent cells (*Escherichia coli* strain DH5 $\alpha$ )
- ❏ LB (Luria Bertani) media
- ❏ Chloramphenicol (stock concentration 25 mg/mL dissolved in EtOH)
- ❏ 50 ml Falcon tube (or equivalent, preferably amber or covered in foil to block light)
- ❏ Incubator at 37°C
- ❏ 1.5 ml eppendorf tubes
- ❏ Ice bucket with ice
- ❏ Micropipettes (capable of pipetting a range of volumes between 1 µL and 1000 µL)
- ❏ Micropipette tips
- ❏ 96 well plates, black with clear flat bottom preferred, at least 3-4 plates (provided by team)

### Calibration Protocols

**CALIBRATION PROTOCOLS SHOULD BE COMPLETED BEFORE CELL MEASUREMENTS ARE TAKEN!**

**You will make three sets of unit calibration measurements: an $OD_{600}$ reference point, a particle standard curve, and a fluorescein standard curve.** Before beginning these protocols, please ensure that you are familiar with the measurement modes and settings of your instrument.

For all of these calibration measurements, you must use the same plates and volumes that you will use in your cell-based assays. You must also use the same settings (e.g., filters or excitation and emission wavelengths) that you will use in your cell-based assays. **If you do not use the same plates, volumes, and settings, the calibration will not be valid.** Make sure to record all information about your instrument (checklist on page 1 of this protocol) as these will be required later when you document your experiment. If your instrument has variable temperature settings, the instrument temperature should be set to room temperature (approximately 20-25 C) for all measurements.

#### Calibration 1: $OD_{600}$ Reference point - LUDOX Protocol

You will use LUDOX CL-X (45% colloidal silica suspension) as a single point reference to obtain a conversion factor to transform your absorbance ($Abs_{600}$) data from your plate reader into a comparable $OD_{600}$ measurement as would be obtained in a spectrophotometer. Such conversion is necessary because plate reader measurements of absorbance are volume dependent; the depth of the fluid in the well defines the path length of the light passing through the sample, which can vary slightly from well to well. In a standard spectrophotometer, the path length is fixed and is defined by the width of the cuvette, which is constant. Therefore this conversion calculation can transform $Abs_{600}$ measurements from a plate reader (i.e., absorbance at 600nm, the basic output of most instruments) into comparable $OD_{600}$ measurements. The LUDOX solution is only weakly scattering and so will give a low absorbance value.

[**IMPORTANT NOTE**: many plate readers have an automatic path length correction feature. This adjustment compromises the accuracy of measurement in highly light scattering solutions, such as dense cultures of cells. **YOU MUST THEREFORE TURN OFF PATHLENGTH CORRECTION if it can be disabled on your instrument**.]

**Materials:**

1ml LUDOX CL-X (provided in kit)
$ddH_20$ (provided by team)
96 well plate, black with clear flat bottom preferred (provided by team)

**Method**

- ❏ Add 100 µl LUDOX into wells A1, B1, C1, D1
- ❏ Add 100 µl of dd $H_2O$ into wells A2, B2, C2, D2
- ❏ Measure absorbance at 600 nm of all samples in the measurement mode you plan to use for cell measurements
- ❏ Record the data in the table below or in your notebook
- ❏ Import data into Excel sheet provided (**OD600 reference point tab**)

| | LUDOX CL-X | $ddH_2O$ |
|---|---|---|
| replicate 1 | | |
| replicate 2 | | |
| replicate 3 | | |
| replicate 4 | | |

| | A | B | C | D |
|---|---|---|---|---|
| 1 | | LUDOX CL-X | H2O | |
| 2 | Replicate 1 | 0.078 | 0.038 | |
| 3 | Replicate 2 | 0.077 | 0.038 | |
| 4 | Replicate 3 | 0.078 | 0.038 | |
| 5 | Replicate 4 | 0.078 | 0.038 | |
| 6 | Arith. Mean | 0.078 | 0.038 | |
| 7 | Corrected Abs600 | 0.040 | | |
| 8 | Reference OD600 | 0.063 | | |
| 9 | OD600/Abs600 | 1.585 | | |
| 10 | | | | |
| 11 | | | | |

The screen capture image above is from the OD600 Reference Point tab of the InterLab Excel sheet. The table shows the data for $OD_{600}$ measured by a spectrophotometer (row 8, yellow box, "Reference OD600") and plate reader data for the $H_2O$ and LUDOX similar to what you will likely collect (you will place your own data in the blue boxes). The corrected $Abs_{600}$ is calculated by subtracting the $H_2O$ reading. The reference $OD_{600}$ is defined as that measured by the reference spectrophotometer (as provided to you in the Excel sheet). The correction factor to convert measured $Abs_{600}$ to $OD_{600}$ is thus the Reference $OD_{600}$ divided by $Abs_{600}$. ***All cell density readings using this instrument with the same settings and volume can be converted to*** $OD_{600}$ ***by multiplying by (in this example) 1.585.***

#### <u>Calibration 2:</u> Particle Standard Curve - Microsphere Protocol

You will prepare a dilution series of monodisperse silica microspheres and measure the $Abs_{600}$ in your plate reader. The size and optical characteristics of these microspheres are similar to cells, and there is a known amount of particles per volume. This measurement will allow you to construct a standard curve of particle concentration which can be used to convert $Abs_{600}$ measurements to an estimated number of cells.

##### Materials:

300 µL Silica beads - Microsphere suspension (provided in kit, 4.7 x 10^8 microspheres)
$ddH_20$ (provided by team)
96 well plate, black with clear flat bottom preferred (provided by team)

##### Method:

**Prepare the Microsphere Stock Solution:**

❑ Obtain the tube labeled "Silica Beads" from the InterLab test kit and vortex vigorously for 30 seconds. **NOTE: Microspheres should NOT be stored at 0°C or below**, as freezing affects the properties of the microspheres. If you believe your microspheres may have been frozen, please contact the iGEM Measurement Committee for a replacement (measurement at igem dot org).

- ❑ Immediately pipet 96 µL microspheres into a 1.5 mL eppendorf tube
- ❑ Add 904 µL of dd$H_2O$ to the microspheres
- ❑ Vortex well. This is your Microsphere Stock Solution.

**Prepare the serial dilution of Microspheres:**

Accurate pipetting is essential. Serial dilutions will be performed across columns 1-11. COLUMN 12 MUST CONTAIN dd$H_2O$ ONLY. Initially you will setup the plate with the microsphere stock solution in column 1 and an equal volume of 1x dd$H_2O$ in columns 2 to 12. You will perform a serial dilution by consecutively transferring 100 µl from column to column with good mixing.

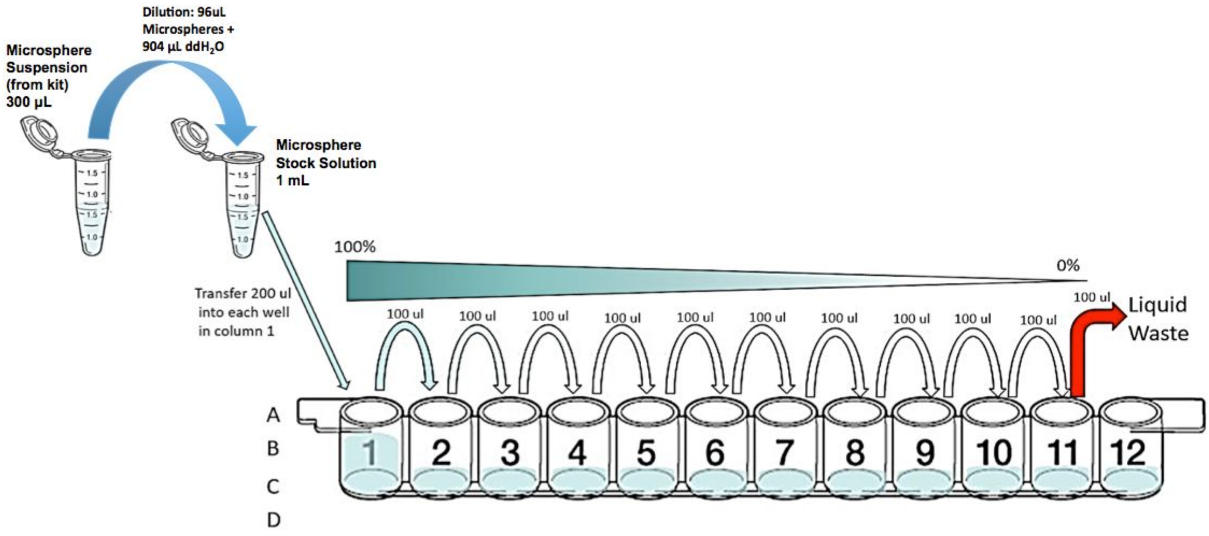


- ❑ **Add 100 µl of dd$H_2O$** into wells A2, B2, C2, D2....A12, B12, C12, D12
- ❑ **Vortex the tube containing the stock solution of microspheres vigorously for 10 seconds**
- ❑ **Immediately add 200 µl of microspheres stock solution into A1**
- ❑ Transfer 100 µl of microsphere stock solution from A1 into A2.
- ❑ Mix A2 by pipetting up and down 3x and transfer 100 µl into A3…
- ❑ Mix A3 by pipetting up and down 3x and transfer 100 µl into A4...
- ❑ Mix A4 by pipetting up and down 3x and transfer 100 µl into A5...
- ❑ Mix A5 by pipetting up and down 3x and transfer 100 µl into A6...
- ❑ Mix A6 by pipetting up and down 3x and transfer 100 µl into A7...
- ❑ Mix A7 by pipetting up and down 3x and transfer 100 µl into A8...

- [ ] Mix A8 by pipetting up and down 3x and transfer 100 µl into A9...
- [ ] Mix A9 by pipetting up and down 3x and transfer 100 µl into A10...
- [ ] Mix A10 by pipetting up and down 3x and transfer 100 µl into A11...
- [ ] Mix A11 by pipetting up and down 3x and transfer 100 µl into **liquid waste**

**TAKE CARE NOT TO CONTINUE SERIAL DILUTION INTO COLUMN 12.**

- [ ] Repeat dilution series for rows B, C, D
- [ ] **IMPORTANT!** Re-Mix (Pipette up and down) each row of your plate ***immediately before*** putting in the plate reader! (This is important because the beads begin to settle to the bottom of the wells within about 10 minutes, which will affect the measurements.) Take care to mix gently and avoid creating bubbles on the surface of the liquid.
- [ ] Measure $Abs_{600}$ of all samples in instrument
- [ ] Record the data in your notebook
- [ ] Import data into Excel sheet provided (**particle standard curve tab**)

#### Calibration 3: Fluorescence standard curve - Fluorescein Protocol

Plate readers report fluorescence values in arbitrary units that vary widely from instrument to instrument. Therefore absolute fluorescence values cannot be directly compared from one instrument to another. In order to compare fluorescence output of test devices between teams, it is necessary for each team to create a standard fluorescence curve. Although distribution of a known concentration of GFP protein would be an ideal way to standardize the amount of GFP fluorescence in our *E. coli* cells, the stability of the protein and the high cost of its purification are problematic. We therefore use the small molecule fluorescein, which has similar excitation and emission properties to GFP, but is cost-effective and easy to prepare. (The version of GFP used in the devices, GFP mut3b, has an excitation maximum at 501 nm and an emission maximum at 511 nm; fluorescein has an excitation maximum at 494 nm and an emission maximum at 525nm).

You will prepare a dilution series of fluorescein in four replicates and measure the fluorescence in a 96 well plate in your plate reader. By measuring these in your plate reader, you will generate a standard curve of fluorescence for fluorescein concentration. You will be able to use this to convert your cell based readings to an equivalent fluorescein concentration. Before beginning this protocol, ensure that you are familiar with the GFP settings and measurement modes of your instrument. You will need to know what filters your instrument has for measuring GFP, including information about the bandpass width (530 nm / 30 nm bandpass, 25-30nm width is recommended), excitation (485 nm is recommended) and emission (520-530 nm is recommended) of this filter.

**Materials:**

Fluorescein (provided in kit)
10ml 1xPBS pH 7.4-7.6 (phosphate buffered saline; provided by team)

96 well plate, black with clear flat bottom (provided by team)

**Method**

**Prepare the fluorescein stock solution:**

- ❑ Spin down fluorescein kit tube to make sure pellet is at the bottom of tube.
- ❑ Prepare 10x fluorescein stock solution (100 µM) by resuspending fluorescein in 1 mL of 1xPBS. [**Note**: it is important that the fluorescein is properly dissolved. To check this, after the resuspension you should pipette up and down and examine the solution in the pipette tip – if any particulates are visible in the pipette tip continue to mix the solution until they disappear.]
- ❑ Dilute the 10x fluorescein stock solution with 1xPBS to make a 1x fluorescein solution with concentration 10 µM: 100 µL of 10x fluorescein stock into 900 µL 1x PBS

**Prepare the serial dilutions of fluorescein:**

Accurate pipetting is essential. Serial dilutions will be performed across columns 1-11. COLUMN 12 MUST CONTAIN PBS BUFFER ONLY. Initially you will setup the plate with the fluorescein stock in column 1 and an equal volume of 1xPBS in columns 2 to 12. You will perform a serial dilution by consecutively transferring 100 µl from column to column with good mixing.

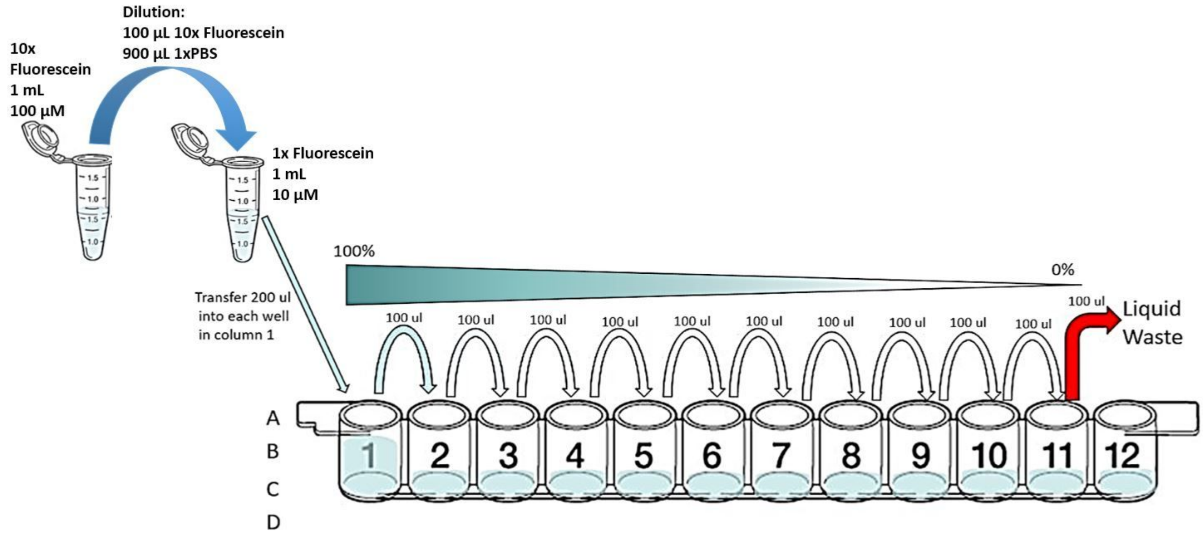


- ❑ **Add 100 µl of PBS** into wells A2, B2, C2, D2....A12, B12, C12, D12
- ❑ **Add 200 µl of fluorescein 1x stock** solution into A1, B1, C1, D1
- ❑ Transfer 100 µl of fluorescein stock solution from A1 into A2.
- ❑ Mix A2 by pipetting up and down 3x and transfer 100 µl into A3…
- ❑ Mix A3 by pipetting up and down 3x and transfer 100 µl into A4...

❑ Mix A4 by pipetting up and down 3x and transfer 100 µl into A5...

❑ Mix A5 by pipetting up and down 3x and transfer 100 µl into A6...

❑ Mix A6 by pipetting up and down 3x and transfer 100 µl into A7...

❑ Mix A7 by pipetting up and down 3x and transfer 100 µl into A8...

❑ Mix A8 by pipetting up and down 3x and transfer 100 µl into A9...

❑ Mix A9 by pipetting up and down 3x and transfer 100 µl into A10...

❑ Mix A10 by pipetting up and down 3x and transfer 100 µl into A11...

❑ Mix A11 by pipetting up and down 3x and transfer 100 µl into **liquid waste**

**TAKE CARE NOT TO CONTINUE SERIAL DILUTION INTO COLUMN 12.**

❑ Repeat dilution series for rows B, C, D

❑ Measure fluorescence of all samples in instrument

❑ Record the data in your notebook

❑ Import data into Excel sheet provided (**fluorescein standard curve tab**)

### Cell measurement protocol

#### Prior to performing the cell measurements you should perform all three of the calibration measurements. Please do not proceed unless you have completed the three calibration protocols.

Completion of the calibrations will ensure that you understand the measurement process and that you can take the cell measurements under the same conditions. For the sake of consistency and reproducibility, we are requiring all teams to use *E. coli* K-12 DH5-alpha. If you do not have access to this strain, you can request streaks of the transformed devices from another team near you, and this can count as a collaboration as long as it is appropriately documented on both teams' wikis. If you are absolutely unable to obtain the DH5-alpha strain, you may still participate in the InterLab study by contacting the Measurement Committee (measurement at igem dot org) to discuss your situation.

For all of these cell measurements, you must use the same plates and volumes that you used in your calibration protocol. You must also use the same settings (e.g., filters or excitation and emission wavelengths) that you used in your calibration measurements. If you do not use the same plates, volumes, and settings, the measurements will not be valid.

**Materials:**

Competent cells (*Escherichia coli* strain DH5 $\alpha$ )
LB (Luria Bertani) media
Chloramphenicol (stock concentration 25 mg/mL dissolved in EtOH)
50 ml Falcon tube (or equivalent, preferably amber or covered in foil to block light)
Incubator at 37°C
1.5 ml eppendorf tubes for sample storage
Ice bucket with ice
Micropipettes and tips
96 well plate, black with clear flat bottom preferred (provided by team)

Devices (from Distribution Kit, all in pSB1C3 backbone):

| **Device** | **Part Number** | **Plate** | **Location** |
|---|---|---|---|
| Negative control | BBa_R0040 | Kit Plate 7 | Well 2D |
| Positive control | BBa_I20270 | Kit Plate 7 | Well 2B |
| Test Device 1 | BBa_J364000 | Kit Plate 7 | Well 2F |
| Test Device 2 | BBa_J364001 | Kit Plate 7 | Well 2H |
| Test Device 3 | BBa_J364002 | Kit Plate 7 | Well 2J |
| Test Device 4 | BBa_J364007 | Kit Plate 7 | Well 2L |
| Test Device 5 | BBa_J364008 | Kit Plate 7 | Well 2N |
| Test Device 6 | BBa_J364009 | Kit Plate 7 | Well 2P |

**Workflow**

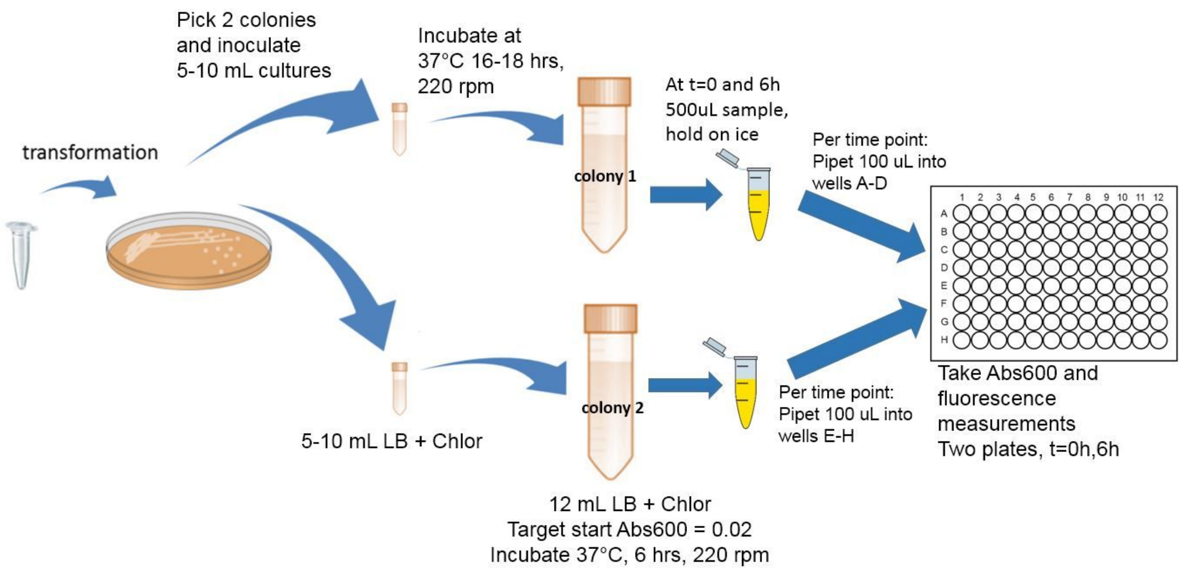


**Method**

**Day 1**: transform *Escherichia coli* DH5 $\alpha$ with these following plasmids (all in pSB1C3):

| Device | Part Number | Plate | Location |
|---|---|---|---|
| Negative control | BBa_R0040 | Kit Plate 7 | Well 2D |
| Positive control | BBa_I20270 | Kit Plate 7 | Well 2B |
| Test Device 1 | BBa_J364000 | Kit Plate 7 | Well 2F |
| Test Device 2 | BBa_J364001 | Kit Plate 7 | Well 2H |
| Test Device 3 | BBa_J364002 | Kit Plate 7 | Well 2J |
| Test Device 4 | BBa_J364007 | Kit Plate 7 | Well 2L |
| Test Device 5 | BBa_J364008 | Kit Plate 7 | Well 2N |
| Test Device 6 | BBa_J364009 | Kit Plate 7 | Well 2P |

**Help Debugging Your Transformations:**

- We STRONGLY recommend that you use the iGEM protocol to create your competent cells: http://parts.igem.org/Help:Protocols/Competent_Cells
- Once you have created your competent cells, we STRONGLY recommend that you measure the competency of your cells using the Competent Cell Test Kit: http://parts.igem.org/Help:2017_Competent_Cell_Test_Kit
- Finally, we STRONGLY recommend that you closely follow the iGEM protocols for resuspending DNA from the kit plates and performing the transformation: http://parts.igem.org/Help:Protocols/Transformation

Year after year, we have found that most teams are highly successful when they follow these protocols, even if alternative protocols are used within your lab. If you are having trouble transforming your test devices, please try the protocols above.

**Day 2**: Pick 2 colonies from each of the transformation plates and inoculate in 5-10 mL LB medium + Chloramphenicol. Grow the cells overnight (16-18 hours) at 37°C and 220 rpm.

**Day 3**: Cell growth, sampling, and assay

- ☐ Make a 1:10 dilution of each overnight culture in LB+Chloramphenicol (0.5mL of culture into 4.5mL of LB+Chlor)
- ☐ Measure $Abs_{600}$ of these 1:10 diluted cultures
- ☐ Record the data in your notebook
- ☐ Dilute the cultures further to a target $Abs_{600}$ of 0.02 in a final volume of **12 ml** LB medium + Chloramphenicol in 50 mL falcon tube (amber, or covered with foil to block light).
- ☐ Take 500 µL samples of the diluted cultures at 0 hours into 1.5 ml eppendorf tubes, prior to incubation. (At each time point 0 hours and 6 hours, you will take a sample from each of the 8 devices, two colonies per device, for a total of 16 eppendorf tubes with 500 µL samples per time point, 32 samples total). Place the samples on ice.
- ☐ Incubate the remainder of the cultures at 37°C and 220 rpm for 6 hours.
- ☐ Take 500 µL samples of the cultures at 6 hours of incubation into 1.5 ml eppendorf tubes. Place samples on ice.
- ☐ At the end of sampling point you need to measure your samples ($Abs_{600}$ and fluorescence measurement), see the below for details.
- ☐ Record data in your notebook
- ☐ Import data into Excel sheet provided (**fluorescence measurement tab**)

**Measurement**

Samples should be laid out according to the plate diagram below. Pipette 100 µl of each sample into each well. From 500 µl samples in a 1.5 ml eppendorf tube, 4 replicate samples of colony #1 should be pipetted into wells in rows A, B, C and D. Replicate samples of colony #2 should be pipetted into wells in rows E, F, G and H. Be sure to include 8 control wells containing 100uL each of only LB+chloramphenicol on each plate in column 9, as shown in the diagram below. Set the instrument settings as those that gave the best results in your calibration curves (no measurements off scale). If necessary you can test more than one of the previously calibrated settings to get the best data (no measurements off scale). Instrument temperature should be set to room temperature (approximately 20-25 C) if your instrument has variable temperature settings.

**Help Debugging:**

- If you have measurements that are off scale (“OVERFLOW”), that data will not be usable. You need to adjust your settings so that the data will be in range and re-run your calibration.
- If your Abs600 measurements for your cell colonies are very close to that of your LB+Chlor, then your cells have probably not been transformed correctly or grown correctly.
- If your negative and positive control values are very close to each other, that probably means something has gone wrong in your protocol or measurement.

**Layout for $Abs_{600}$ and Fluorescence measurement**

At the end of the experiment, you should have two plates to read. Each plate should be set up as shown below. You will have one plate for each time point: 0 and 6 hours. On each plate you will read both fluorescence and absorbance.

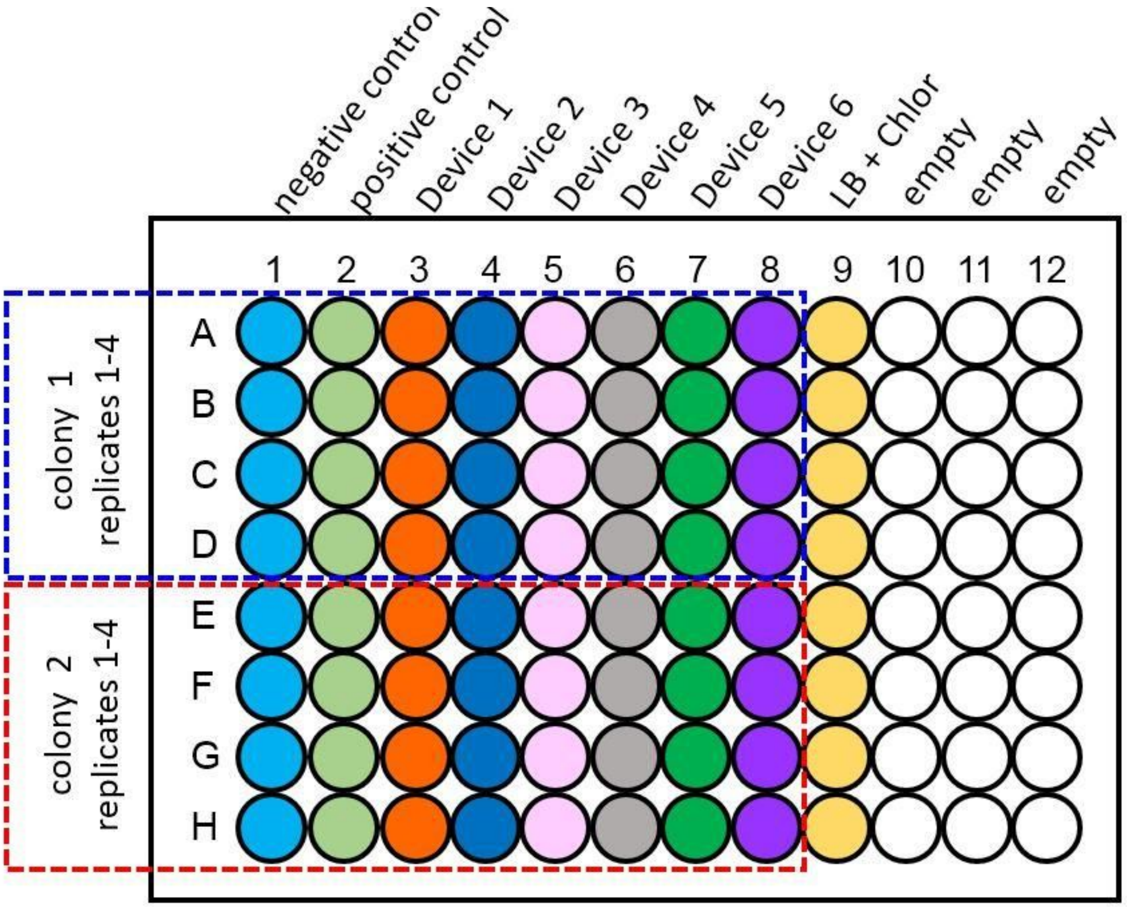

### Protocol: Colony Forming Units per 0.1 $OD_{600}$ *E. coli* cultures

This procedure can be used to calibrate $OD_{600}$ to colony forming unit (CFU) counts, which are directly relatable to the cell concentration of the culture, i.e. viable cell counts per mL. This protocol assumes that 1 bacterial cell will give rise to 1 colony.

For the CFU protocol, you will need to count colonies for your two Positive Control (BBa_I20270) cultures and your two Negative Control (BBa_R0040) cultures.

**Step 1: Starting Sample Preparation**
This protocol will result in CFU/mL for 0.1 $OD_{600}$. Your overnight cultures will have a much higher $OD_{600}$ and so this section of the protocol, called "Starting Sample Preparation", will give you the "Starting Sample" with a 0.1 $OD_{600}$ measurement.

1. Measure the $OD_{600}$ of your cell cultures, making sure to dilute to the linear detection range of your plate reader, e.g. to 0.05 – 0.5 $OD_{600}$ range. Include blank media (LB + Cam) as well.

   For an overnight culture (16-18 hours of growth), we recommend diluting your culture 1:8 (8-fold dilution) in LB + Cam before measuring the $OD_{600}$.

   **Preparation:** Add 25 µL culture to 175 µL LB + Cam in a well in a black 96-well plate, with a clear, flat bottom.

   Recommended plate setup is below. Each well should have 200 µL .

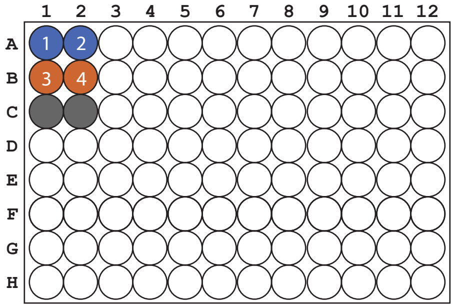


A1 - A2: Positive Controls (cultures 1-2)
B1 - B2: Negative Controls (cultures 3-4)
C1 - C2: Blank media - 200 µL of LB + Cam (in duplicate)

2. Dilute your overnight culture to $OD_{600}$ = 0.1 in 1mL of LB + Cam media. Do this in triplicate for each culture.

   Use $(C_1)(V_1) = (C_2)(V_2)$ to calculate your dilutions

   | | |
   |---|---|
   | $C_1$ is your starting $OD_{600}$ | $C_2$ is your target $OD_{600}$ of 0.1 |
   | $V_1$ is the unknown volume in µL | $V_2$ is the final volume of 1000 µL |

   **Important:** When calculating $C_1$, subtract the blank from your reading and multiple by the dilution factor you used.
   Example: C1 = (1:8 $OD_{600}$ - blank $OD_{600}$) x 8 = (0.195 - 0.042) x 8 = 0.153 x 8 = 1.224

   Example: $(C_1)(V_1) = (C_2)(V_2)$
   (1.224)(x) = (0.1)(1000µL)
   x = 100/1.224 = 82 µL culture
   Add 82 µL of culture to 918 µL media for a total volume of 1000 µL

3. Check the $OD_{600}$ and make sure it is 0.1 (minus the blank measurement).
   Recommended plate setup is below. Each well should have 200 µL .

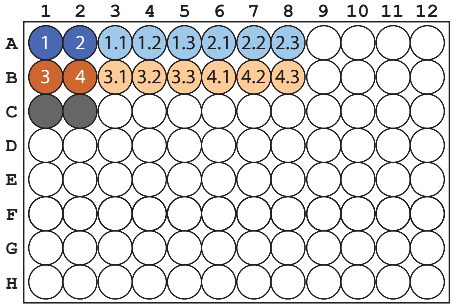


A1 - A2: Positive Controls (cultures 1-2)
B1 - B2: Negative Controls (cultures 3-4)
C1 - C2: Blank media - 200 µL of LB + Cam (in duplicate)
A3 - A8: 0.1 Starting Sample Dilutions for Positive Controls (in triplicate per culture, 6 total dilutions)
B3 - B8: 0.1 Starting Samples Dilutions for Negative Controls (in triplicate per culture, 6 total dilutions)

**Step 2: Dilution Series Instructions**

Do the following serial dilutions for your triplicate Starting Samples you prepared in Step 1. You should have 12 total Starting Samples - 6 for your Positive Controls and 6 for your Negative Controls.

For each Starting Sample (*total for all 12 showed in italics in paraenthesis*):

1. You will need 3 LB Agar + Cam plates (*36 total*).
2. Prepare three 2.0 mL tubes (*36 total*) with 1900 µL of LB + Cam media for Dilutions 1, 2, and 3 (see figure below).
3. Prepare two 1.5 mL tubes (*24 total*) with 900 µL of LB + Cam media for Dilutions 4 and 5 (see figure below)..
4. Label each tube according to the figure below (Dilution 1, etc.) for each Starting Sample.
5. Pipet 100 µL of Starting Culture into Dilution 1. Discard tip. Do NOT pipette up and down. Vortex tube for 5-10 secs.
6. Repeat Step 5 for each dilution through to Dilution 5 as shown below.
7. Aseptically spead plate 100 µL on LB + Cam plates for Dilutions 3, 4, and 5.
8. Incubate at 37°C overnight and count colonies after 18-20 hours of growth.

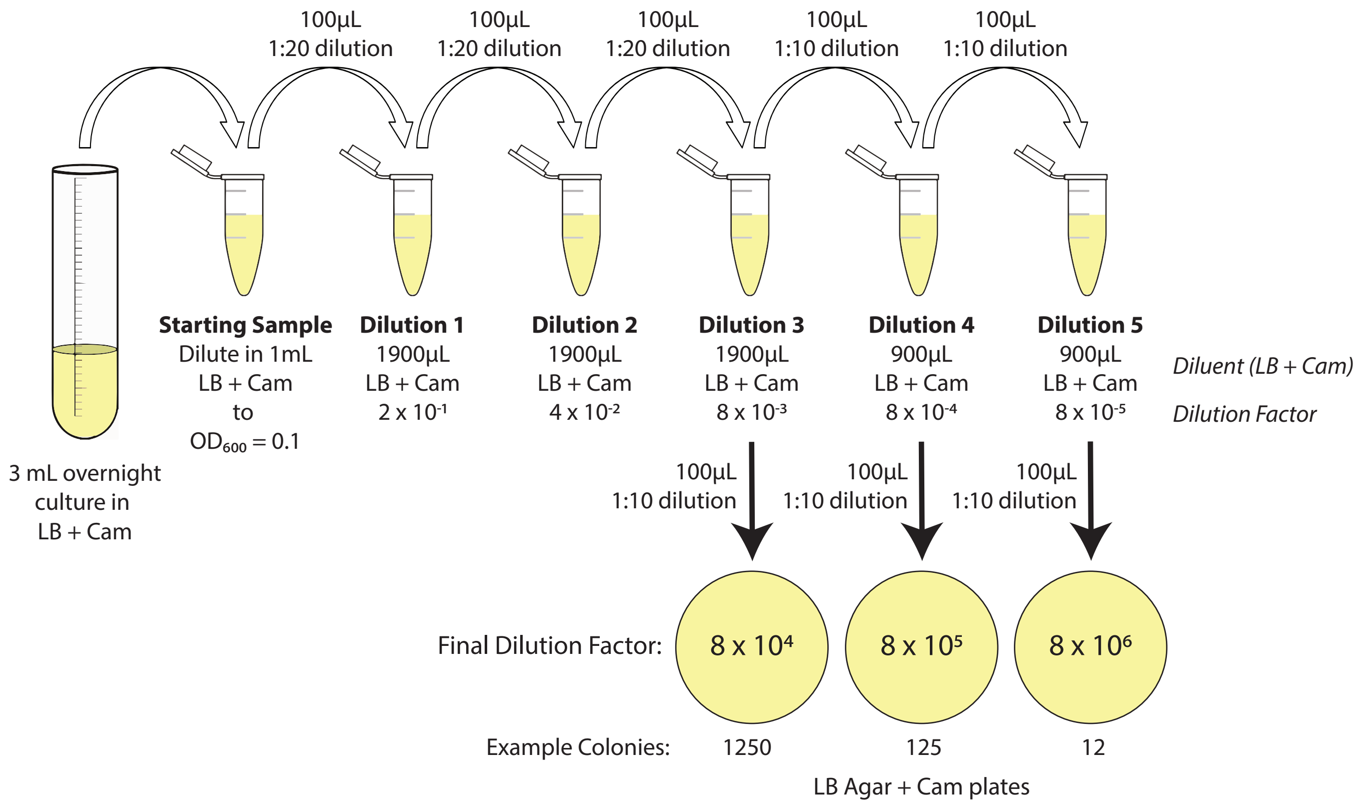


**Step 3: CFU/mL/OD Calculation Instructions**
Based on the assumption that 1 bacterial cell gives rise to 1 colony, colony forming units (CFU) per 1mL of an $OD_{600}$ = 0.1 culture can be calculated as follows:

1. Count the colonies on each plate with fewer than 300 colonies.
2. Multiple the colony count by the Final Dilution Factor on each plate.

Example using Dilution 4 from above

| | | | |
|---|---|---|---|
| # colonies | x | Final Dilution Factor | = CFU/mL |
| 125 | x | $(8 \times 10^5)$ | = $1 \times 10^8$ CFU/mL in Starting Sample ($OD_{600}$ = 0.1) |
