## Supplementary Note: Flow Cytometer Protocol for "Robust Estimation of Bacterial Cell Count from Optical Density"

### Extra Credit: Flow Cytometry

For extra credit, teams with access to a flow cytometer and SpheroTech calibration beads can collect and submit flow cytometry data as well. Teams performing this additional measurement will be given special acknowledgement at the iGEM Jamboree and in any resulting scientific publications.

**Materials:**
SpheroTech Rainbow calibration beads, type RCP-30-5A or URCP-38-2K
([http://www.spherotech.com/CalibrationParticles.htm](http://www.spherotech.com/CalibrationParticles.htm))

Record the lot number for your calibration beads.
It should be one or two letters followed by a number (e.g., “AJ02”)

**Method:**
During your cell measurement protocol, prepare a sample of SpheroTech beads according to the manufacturer instructions and place in well A10 of each plate.

**Measurement:**
After measuring each plate with your plate reader, also collect data from all wells using your flow cytometer. Follow your flow cytometer instructions for collecting samples and dilute further if necessary. Collect at least 10,000 events per well.

On the interlab form, mark that you have done the flow cytometry extra credit, and enter your instrument information in the fields provided.

Name the FCS files for your experimental samples following these templates:

- Cell samples: [team]_[time]h_[well]_[construct].fcs
  *example: WPI_6h_A1_NegativeControl.fcs*
- Blanks: [team]_[time]h_[well]_Blank.fcs
  *example: WPI_6h_A9_Blank.fcs*
- Beads: [team]_[time]h_[well]_[type]_[lot].fcs
  *example: WPI_6h_A10_URCP-38-2K_AJ02.fcs*

Bundle all FCS files together into a zip or tar file and upload to DropBox at:
http://2018.igem.org/Measurement/InterLab/Flow_Cytometry

If you cannot access DropBox, email the measurement committee to make alternate arrangements for delivering your files.
