## Supplementary Note: Data Acceptance Criteria for "Robust Estimation of Bacterial Cell Count from Optical Density"

### S5 File Data Acceptance Criteria

Each data set used in the interlab study was evaluated against the following criteria to determine whether it was of sufficient quality for inclusion. These criteria were not intended to be stringent, but rather to represent a minimal "sanity check" against major errors in protocol execution or reporting. Any teams whose data did not meet all of these criteria were invited to re-execute the experiment in order to correct the deficiencies in their data.

The criteria are:

- Water measurements have a lower OD than LUDOX measurements.
- Silica microsphere OD measurements generally decrease with increasing dilution (excepting saturation)
- Water OD measurements are not negative for either the LUDOX or silica microsphere protocols.
- Fluorescein fluorescence measurements generally decreases with increasing dilution (excepting saturation)
- PBS-only fluorescence measurements are not negative.
- Cell sample fluorescence and OD measurements are within the range covered by silica microsphere and fluorescein samples.
- Positive control is brighter than negative control at 6 hours and also greater than zero
- At least half of cell sample ODs increase significantly from 0 hours to 6 hours (i.e., cells are generally alive and growing)
- Fluorescence/OD measurements for test constructs at 6 hours span at least a 10-fold range (i.e., there is at least some significant variability in fluorescence expression)
- All replicates are present for every sample.
