## Supplementary figures and images for "Robust Estimation of Bacterial Cell Count from Optical Density"

### Supplementary Figure 1 Length of Valid Sequence

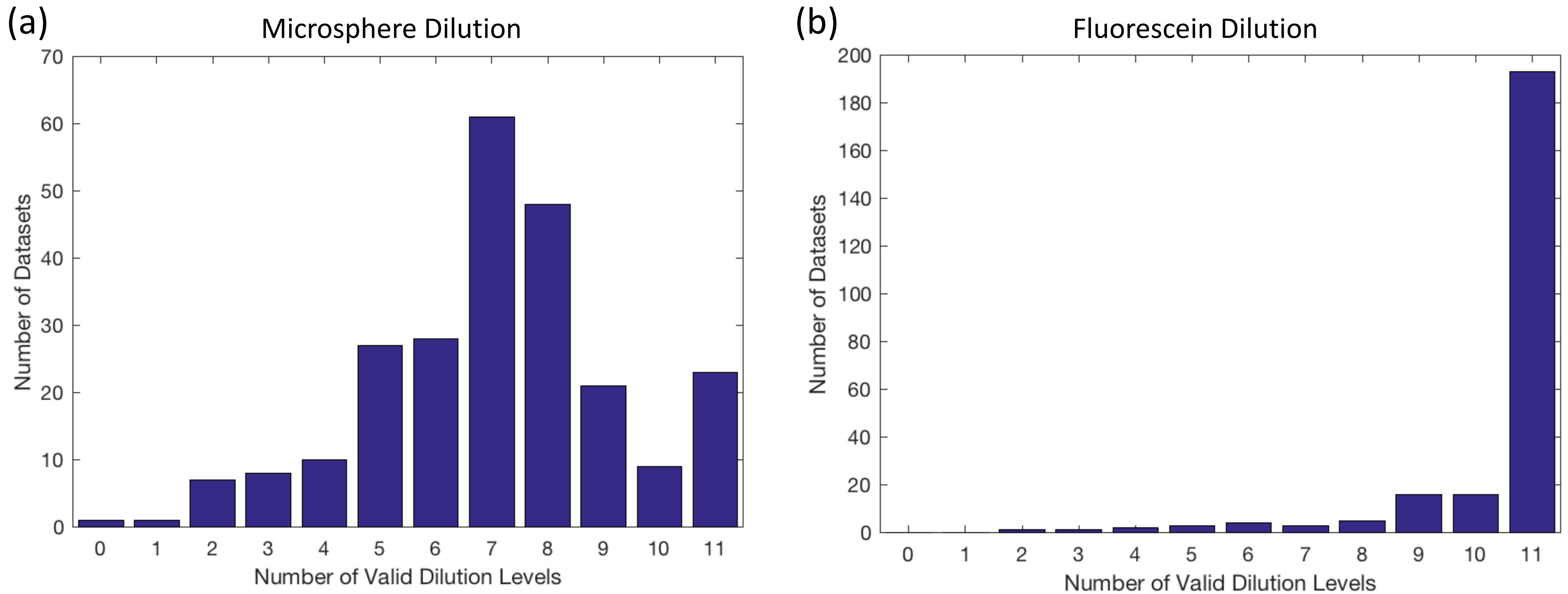

### Supplementary Figure 2 E. coli Colony Growth

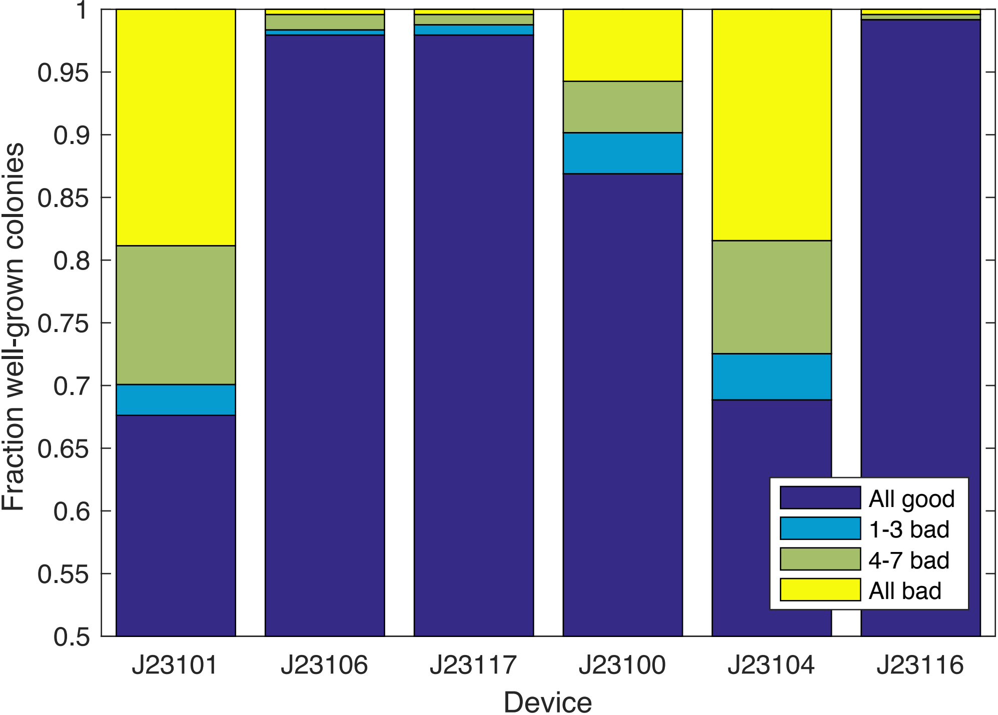

### Supplementary Figure 3 Example of Flow Cytometry Gating

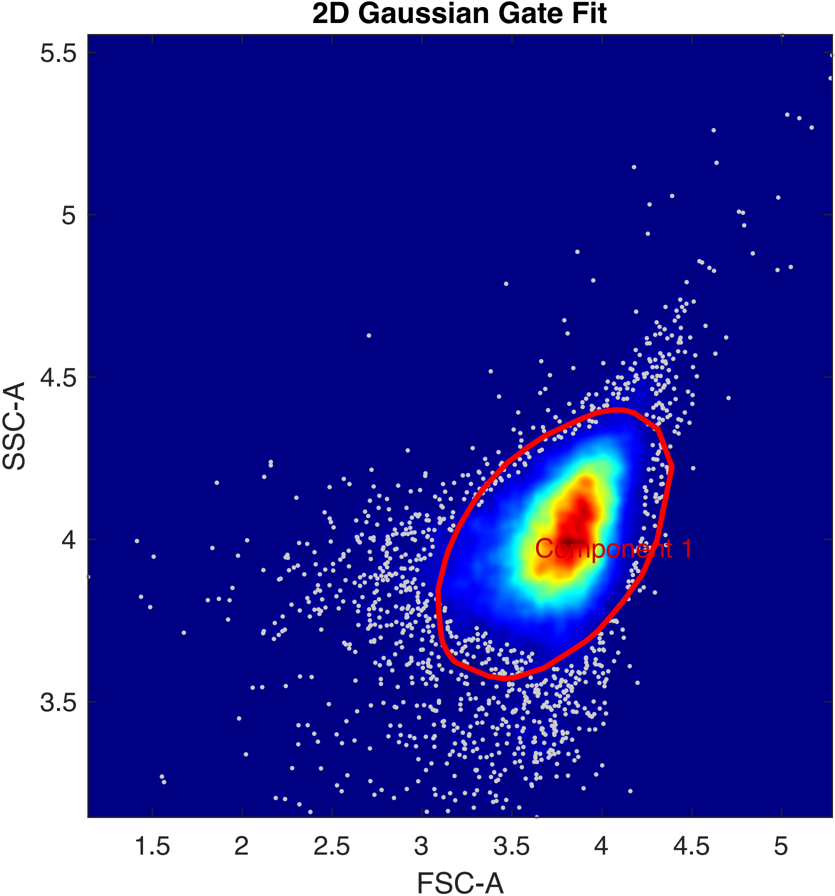
